# Amniotic Fluid Reduces Liver Fibrosis By Attenuating Hepatic Stellate Cell Activation

**DOI:** 10.1101/2025.02.20.639215

**Authors:** Charles M. Bowen, Frederick Ditmars, Naiyou Liu, Jose Marri Abril, David Ajasin, William K. Russell, Heather L. Stevenson, Eliseo A. Eugenin, Jeffrey H. Fair, W. Samuel Fagg

**Affiliations:** Division of Transplant, Department of Surgery, University of Texas Medical Branch, Galveston, Texas, 77555, USA; John Sealy School of Medicine, University of Texas Medical Branch, Galveston, Texas, 77555, USA; Department of Obstetrics and Gynecology, University of Texas Medical Branch, Galveston, Texas, 77555, USA; Department of Neurobiology, Cell Biology, and Anatomy, University of Texas Medical Branch, Galveston, Texas, 77555, USA; Department of Pathology, University of Texas Medical Branch, Galveston, Texas, 77555, USA; Deparment of Biochemistry and Molecular Biology, University of Texas Medical Branch, Galveston, Texas, 77555, USA; Merakris Therapeutics, RTP Frontier, Research Triangle Park, North Carolina, 27709, USA

**Keywords:** Liver fibrosis, regenerative medicine, amniotic fluid, hepatic stellate cell, myofibroblast activation, epithelial-mesenchymal transition

## Abstract

Regardless of the source of injury or metabolic dysfunction, fibrosis is a frequent driver of liver pathology. Excessive liver fibrosis is caused by persistent activation of hepatic stellate cells (HSCs), which is defined by myofibroblast activation (MFA) and the epithelial-mesenchymal transition (EMT). Strategies to prevent or reverse this HSC phenotype will be critical for successful treatment of liver fibrosis. We have previously shown that full-term, cell-free human amniotic fluid (cfAF) inhibits MFA and EMT in fibroblasts *in vitro*. We hypothesize that cfAF treatment can attenuate HSC activation and limit liver fibrosis. We tested if cfAF could prevent liver fibrosis or HSC activation in murine models of liver damage, three-dimensional hepatic spheroids, and HSC cultures. Administering cfAF prevented weight loss and the extent of fibrosis in mice with chronic liver damage without stimulating deleterious immune responses. Gene expression profiling and immunostaining indicated that cfAF administration in carbon tetrachloride-treated mice reduced EMT- and MFA-related biomarker abundance and modulated transcript levels associated with liver metabolism, immune regulatory pathways, and cell signaling. cfAF treatment lowered MFA biomarker levels in a dose-dependent manner in *ex vivo* hepatic spheroids. Treating HSCs with cfAF *in vitro* strongly repressed EMT. Multi-omics analyses revealed that it also attenuates TGFβ-induced MFA and inflammation-associated processes. Thus, cfAF treatment prevents liver fibrosis by safeguarding against persistent HSC activation. These findings suggest that cfAF may be a safe and effective therapy for reducing liver fibrosis and preventing the development of cirrhosis and/or hepatocellular carcinoma.

**Significance:** Chronic activation of hepatic stellate cells leads to irreversible liver fibrosis, and durable treatment options for liver fibrosis remain suboptimal, creating a critical clinical need. Amniotic fluid (AF) reduces inflammation and promotes tissue remodeling and regeneration when applied to cutaneous wounds and in models thereof. However, no prior studies have explored the anti-fibrotic benefits of AF in liver disease. Data from this study indicate that AF can repress liver fibrosis by reducing pro-fibrotic hepatic stellate cell myofibroblast activation. Thus, these findings lay the foundation for early-phase clinical trials investigating cell-free AF as a therapeutic treatment for liver fibrosis and as a bridging therapy for liver transplantation.

## Introduction

Chronic liver disease is the 11^th^ leading cause of death in the United States^1^ and places a significant strain on annual healthcare costs^2,3^. It can arise from various etiologies such as viral infections, alcoholic liver disease, autoimmune hepatitis, and non-alcoholic steatohepatitis^4^. While its canonical progression occurs along a spectrum in which early fibrosis is reversible, this stage is often clinically unreported. This changes upon progression to fulminant liver disease, at which point liver transplantation is the only curative option^5,6,7^. Unfortunately, there is a lack of effective therapeutic interventions to prevent and/or reverse liver fibrosis^8^. Therefore, novel treatments that can safely and effectively inhibit or reverse liver fibrosis are urgently needed.

The underlying pathophysiology of liver fibrosis is complex; how it develops depends on the insult’s source, extent, and persistence. Regardless of its origin, damaged hepatocytes release pro-inflammatory factors that disrupt the cellular homeostasis of neighboring hepatocytes and other liver non-parenchymal cells, including cholangiocytes, endothelial cells, immune cells, and hepatic stellate cells (HSCs)^9^. While this causes cell type-specific alterations, a common theme amongst them is increased activity and abundance of TGFβ, IL-1a, hedgehog ligands, and mesenchymal potentiators, including SLUG and SNAIL^4^. This stimulates normally quiescent HSCs to transdifferentiate into myofibroblasts, proliferate, and undergo the epithelial-mesenchymal transition (EMT)^10–12^. Interestingly, one defining factor of the myofibroblast activation (MFA) phenotype of HSCs is the excessive secretion of extracellular matrix (ECM) proteins^13^. This physically makes up the fibrotic lesion in the liver^4,6^. Therefore, MFA HSCs are the effector cells of liver fibrosis, and strategies to prevent or reverse their activation could constitute viable therapeutic options.

The use of full-term human amniotic fluid (AF) for therapeutic purposes dates back a century^14^. Given that AF supports fetal development throughout gestation, it is unsurprising that it contains immunomodulatory chemokines, pro-growth and anti-inflammatory cytokines, and other regenerative biomolecules^15–19^. Its most widespread and successful clinical use has been in wound healing^20^. It has also been used to treat orthopedic dysfunction^21–24^ and COVID-19 patients with acute respiratory distress syndrome^25,26^. Cell-free AF (cfAF) administration has been safe in humans across these various real-world treatment scenarios, irrespective of the delivery method. It remains unknown, however, what other applications cfAF might have in human health and disease and how its therapeutic effects are conveyed at the biochemical, molecular, or cellular levels. We began to address the latter question and discovered that cfAF represses EMT and MFA in fibroblasts and myoblasts cultured *in vitro*^27^. Given that HSC transdifferentiation into pro-inflammatory activated HSCs is mediated through MFA and EMT, we hypothesized that cfAF might prevent liver fibrosis by attenuating HSC activation.

We test this using mouse models of liver fibrosis, a human hepatic spheroid model, and two-dimensional cell culture of human hepatic stellate cells. Administering human cfAF to mice is safe and reduces liver fibrosis in the carbon tetrachloride (CCl_4_) model of chronic liver fibrosis and an ethanol-treated human hepatic spheroid model. Transcriptome analysis of CCl_4_-treated mice reveals that cfAF’s protective effects are mediated through the downregulation of MFA, EMT, and signaling pathways, including Wnt, TGFβ, MAPK, mTOR, and NF-κB. Additionally, cfAF treatment prevents cell migration and EMT in human HSCs *in vitro*. Multi-omics analyses reveal that cfAF attenuates EMT- and MFA-associated gene expression and signaling pathways, including those associated with TGFβ, Wnt, Hedgehog, TNF/NFkB, mTOR, and MAPK. These findings demonstrate that cfAF can safely prevent liver fibrosis across a xenogeneic species mismatch. This protective effect is mediated, at least in part, by repressing HSC activation. This study thus suggests that cfAF has the potential to safely and effectively treat patients with fibrotic liver disease.

## Materials and Methods

### Ethics statement

All patient samples (cfAF) and information were collected with written informed consent. Animal care and husbandry conformed to practices established by the Association for the Assessment and Accreditation of Laboratory Animal Care (AAALAC), The Guide for the Care and Use of Laboratory Animals, and the Animal Welfare Act. The Institutional Animal Care and Use Committee (IACUC) of The University of Texas Medical Branch approved all animal experiments in accordance with institutional guidelines (IACUC protocol #1706040B).

### Animal Experiments

Wild-type female C57BL/6J mice between 6-8 weeks of age were purchased from Jackson Laboratory (Strain #:000664, Bar Harbor, ME). Mice were grouped based on test condition: sham control (n = 10), cfAF only (n = 7), CCl_4_ only (n = 13), or CCl_4_ plus cfAF (n = 12), for the CCl_4_ chronic liver fibrosis study; and sham control (n = 5), DMN only (n = 5), or DMN plus cfAF (n = 5) for the DMN acute liver damage study. Mice were treated twice weekly for nine weeks with CCl_4_ (2.25 µL/gram body weight or 10 mg/kg DMN) diluted 1:3 in food-grade olive oil or sham control (olive oil only). Either cfAF or saline control was injected i.p. twice weekly for nine weeks (on days when mice did not receive either DMN or CCl_4_) at a dose of 12.5 µL/gram body weight (stock cfAF [1 mg/mL]) or comparable volume of saline. Blood was collected using a capillary tube via the retroorbital approach under anesthesia at baseline, or at the terminal endpoint via direct cardiac puncture. Mice were weighed weekly to monitor health and fitness.

Please see Supplemental Materials and Methods section for additional information.

## Results

### Administration of cfAF in mice is safe and prevents failure to thrive in the carbon tetrachloride liver fibrosis model

We hypothesized that treating wild-type C57BL/6J mice with human cfAF would prevent liver fibrosis without triggering a harmful immune response, as it is immune-privileged and devoid of cells^14,41^. We tested this hypothesis using CCl_4_ to model chronic liver damage,^42–44^ and treated with or without cfAF **(Figure 1A)**. Briefly, mice were treated twice weekly for nine weeks with either CCl_ (2.25 µL/g body weight) diluted 1:3 in food-grade olive oil, or with sham control (olive oil only). On alternating days, mice received i.p. injection of either cfAF (12.5 µL/g body weight of 1 mg/mL stock) or an equivalent volume of saline, also twice weekly for nine weeks. The successful induction of this well-characterized model of liver fibrosis is confirmed by the presence of elevated liver proteins in the serum, failure to thrive (reduced weight gain in 10-week-old mice compared to controls), and histological analysis^42–44^. We detected elevated serum ALT levels in mice treated with CCl_4_ (with or without cfAF) compared to wild-type controls **(Suppl Figure 1A**). No significant differences in serum AST levels were observed between treatment groups (One-way ANOVA; **Suppl Figure 1A**). Mice treated with CCl_4_ alone did not gain weight as quickly or to the same extent as the control mice over the course of the experiment (terminal *P* < 0.0001 by two-way ANOVA; **Figure 1B**). Interestingly, administering cfAF to CCl_4_ mice prevented inefficient weight gain, effectively rescuing the failure-to-thrive phenotype (**Figure 1B; Suppl Figure 1B**).

**Figure 1.**
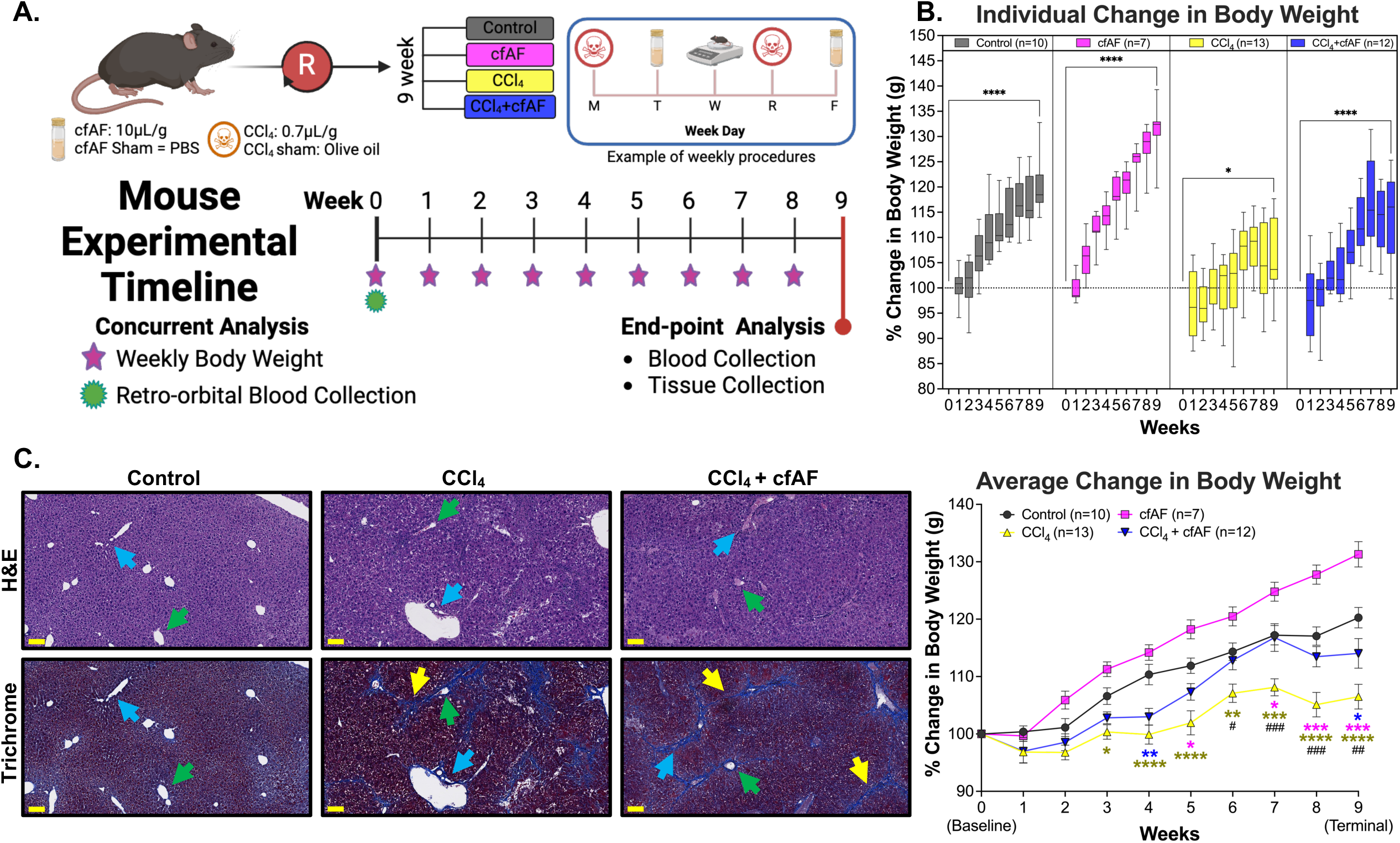
Cell-free amniotic fluid (cfAF) administration rescues failure-to-thrive in carbon tetrachloride (CCl_4_)- treated mice. **A.** Overview of experimental design for modeling chronic liver damage and cfAF administration in mice. **B**. Mean percent weight change by week, relative to starting body weight plotted by treatment (top) or comparing amongst treatment groups (bottom) (*^/#^*P* < 0.05, **^/##^*P* < 0.01, ***^/###^*P* < 0.001, \*\*\*\**P* < 0.0001 by Two-way ANOVA with multi-comparisons; *indicates comparisons between experimental groups and the control, while #represents comparisons between CCl4 + cAF mice and CCl4 mice). **C.** Liver histology using H&E or Masson’s trichrome stains of matched samples showing approximately the same field of view; scale bars = 100um; Blue arrows indicate a portal triad, green arrows indicate the hepatic vein, and yellow arrows mark areas of fibrosis.

We also observed liver fibrosis and altered architecture of the portal tracts in CCl_4_-treated mice (**Figure 1C**), as expected^42,45^. A blinded histological analysis of H&E-stained livers from control, CCl_4_-treated, and CCl_4_-treated + cfAF mice did not show any significant differences in each damage-associated metric, although there was a slight reduction in apoptosis, perivenular inflammation, and ceroid-laden macrophages in the CCl_4_ + cfAF mice compared to CCl_4_-only mice (**Figure 1C**; **Suppl Figure 1C**). Staining liver tissue with Masson’s Trichrome indicated extensive peri-portal (*blue arrows* show portal tracts) fibrosis (*yellow arrows*) bridging to the central vein (*green arrows*) in CCl_4_-treated mice (**Figure 1C; Suppl Figure 1C**). We observed a moderate decrease in fibrosis in the CCl_4_ + cfAF mice, while no evidence of fibrosis was seen in control mice (**Figure 1C**).

To characterize the immunological changes that might be elicited by cfAF administration, we profiled cells collected from the spleen and peripheral blood mononuclear cells (PBMCs) collected from whole blood. Next, we measured immunologic markers for T-cells (CD3, CD4, CD8), B-cells (CD45R/B220), neutrophils (Ly6G^+^/Ly6C^-^), and macrophages (Ly6G^-^/Ly6C^+^). Flow cytometry analysis showed there were no significant differences in the immune cell distribution of PBMCs (**Suppl Figure 1D, left**) or viable splenocyte abundance (**Suppl Figure 1D, right**) between the groups.

In addition to the CCl_4_ model of chronic liver fibrosis, similar experiments were conducted using the dimethylnitrosamine (DMN) murine model of acute liver damage (**Suppl Figure 2A)**^46,47^. Mice treated with DMN, with or without cfAF, experienced significant weight loss over the three-week study period compared to control mice (Terminal *P* < 0.0001 for +DMN and 0.003 for +DMN +cfAF by two-way ANOVA; **Suppl Figure 2B**). Mice treated with DMN alone had elevated serum AST activity compared to control mice as expected, and mice treated with DMN and cfAF had slightly lower levels compared to DMN alone (*P* = 0.6258 by Student’s t-test; **Suppl Figure 2C, right**). There was no significant difference in serum ALT levels between treatment groups; however, serum levels were lowest in the control mice, followed by the DMN + cfAF mice, and highest in the DMN-only treated mice (**Suppl Figure 2C, left**). Masson’s trichrome staining and histological analysis indicated that DMN-treated mice showed evidence of fibrosis, regardless of cfAF administration (**Suppl Figure 2D**). Thus, cfAF administration to immune-competent mice is safe and well-tolerated, and limits failure-to-thrive and liver fibrosis in the CCl4 mouse model of chronic liver damage.

### Cell-free amniotic fluid treatment reduces CCl_4_-induced liver fibrosis and attenuates hepatic stellate cell activation

We utilized the Nanostring platform to measure transcriptome-level changes in 770 fibrosis-associated genes from RNA extracted from the mouse livers mentioned above (**Figure 2A**). We first investigated how similar the transcriptome profiles were among the control, CCl_4_, and CCl_4_ + cfAF mice. Clustering analysis indicates that the control mice exhibited the most dissimilar profile (**Figure 2B, Suppl Table S1**). Aside from a single outlier, the profiles of CCl_4_ + cfAF-treated mice clustered more closely to the control samples than the CCl_4_-treated mice clustered with control, indicating a more similar fibrosis-associated transcriptome to control than in CCl_4_ only compared to control (**Figure 2B**). Analysis of the overlap between each group revealed that 169 significantly altered transcripts were shared between all three, with 29 altered only in the CCl_4_ group compared to control, and 45 altered solely in the CCl_4_ + cfAF group compared to control (*P* < 0.1; **Figure 2C**). When comparing the CCl_4_ + cfAF to the CCl_4_ group, 41 uniquely altered transcripts were identified, likely due to the cfAF treatment (**Figure 2C**). The distribution of transcripts altered under CCl4 + cfAF administration compared to CCl_4_ treatment alone showed increases from certain immune-related genes (*Adcy7, H2-DMa, Bst2*) and decreases from inflammatory-associated and cell cycle regulatory genes (*Kras, Jak1, Map3k1, Acsl4*; **Figure 2D**). This led us to perform pathway analysis, which revealed altered enrichment of 51 fibrosis-associated pathways (**Suppl Figure 3)**^48–50^. Notably, after treating CCl_4_ mice with cfAF, we observed a reduction in the abundance of transcripts linked to EMT, MFA, angiogenesis, and Wnt signaling compared to CCl_4_-treated mice, alongside increases in M1 macrophage, type I and II interferon, and Th1 differentiation (**Figure 2E**). We validated the results obtained from the Nanostring analysis using RT-qPCR to measure the mRNA abundance of Snail, Ncad, and Vim (associated with EMT) and Col1a1 (associated with MFA). This corroborated our Nanostring findings and shows reduced levels of EMT and MFA biomarkers in cfAF-treated mice compared to those receiving only CCl_4_ treatment (**Figure 2F**). Collectively, these results indicate that cfAF treatment prevents CCl_4_-induced liver fibrosis, partly by attenuating HSC activation.

**Figure 2.**
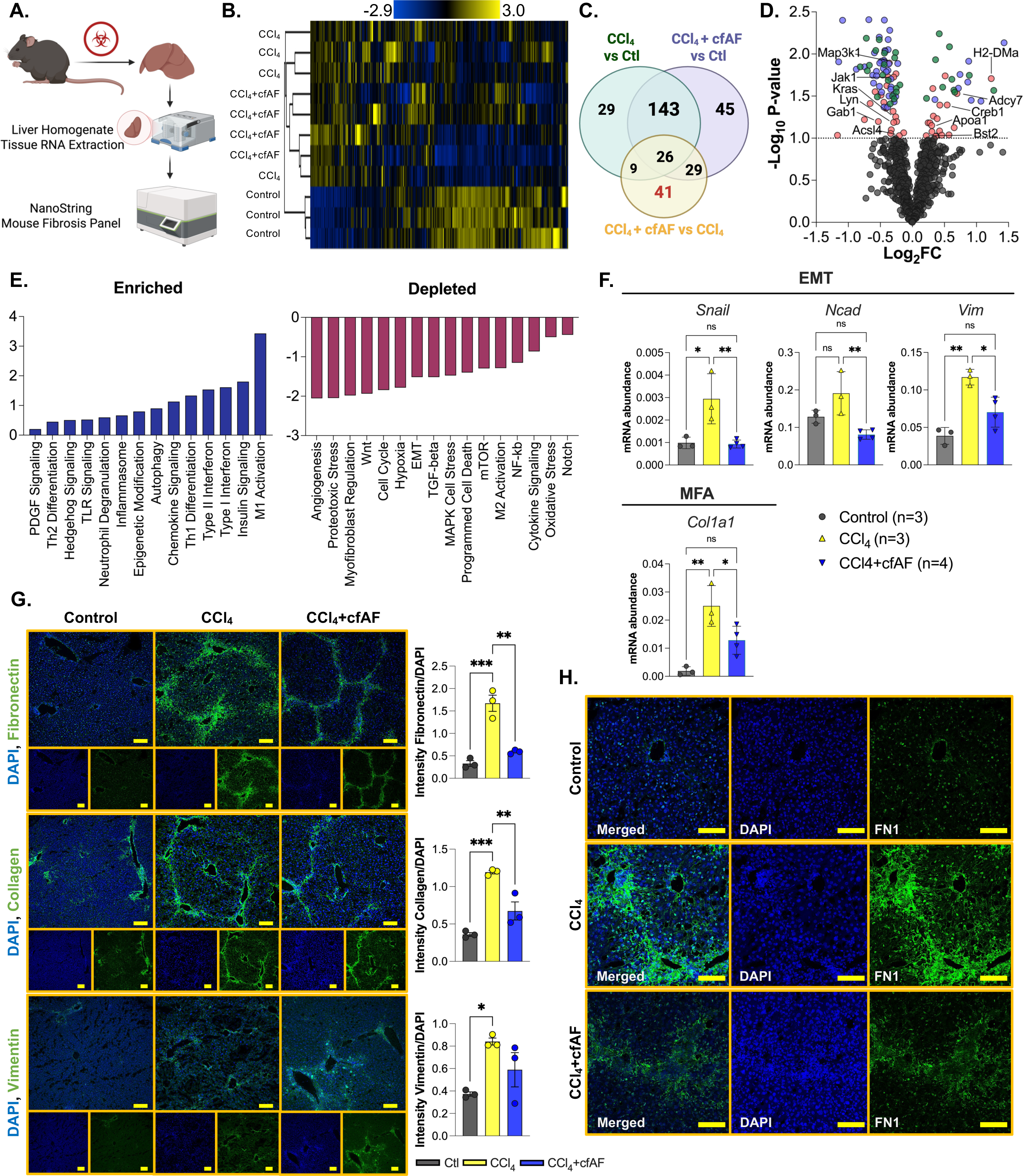
cfAF treatment reduces CCl_4_-induced liver fibrosis by attenuating hepatic stellate cell activation. **A.** Experimental approach showing how mouse liver RNA was analyzed using the Nanostring nCounter Mouse Fibrosis Panel. **B.** Clustered heatmap showing RNA abundance from Nanostring nCounter normalized read levels clustered using Spearman’s rank correlation and single linkage. **C.** A Venn diagram showing the overlap of significant RNA abundance changes (*P* < 0.1) in each comparison from Nanostring nCounter results. **D.** Nanostring data shown in volcano plot with Log_2_FC on the x-axis and -Log_10_ *P*-value on the y-axis; *P*-value cutoff for significance is *P* < 0.1 shows transcripts altered between CCl_4_ vs. control (green), CCl_4_ + cfAF vs. control (blue), and CCl_4_ + cfAF vs. CCl_4_ (red). **E.** Pathway analysis of RNA level changes associated with enrichment (left) or depletion (right) of pathways in CCl_4_ + cfAF mice versus CCl_4_ mice. **F.** RT-qPCR using RNA extracted from murine liver tissue and showing mRNA abundance relative to GAPDH. One-way ANOVA with multi-comparison was used to determine significance: \**P* < 0.05, \*\**P* < 0.01; N = 3 CCl_4_ mice, N = 4 CCl_4_ + cfAF mice; N = 3 control mice. **G.** Representative immunofluorescent images from murine liver tissue using antibodies to Fibronectin (top), Type I Collagen (middle), or Vimentin (bottom) in green and DAPI counterstain in blue; antibody signal intensity is shown relative to DAPI and displayed to the right of each set of images; N=3 mice per treatment condition; scale bars = 100 µm; One-way ANOVA with multi-comparison was used to determine significance: \**P* < 0.05, \*\**P* < 0.01, \*\*\**P* < 0.001. **H.** Higher magnification immunofluorescent images showing Fibronectin antibody and DAPI counterstain focused on an individual portal triad; scale bars = 100µm.

To further evaluate whether cfAF treatment inhibits HSC activation, we employed antibody staining and immunofluorescence (IF) analysis of liver tissue sections to assess the *in situ* protein abundance of biomarkers for EMT and MFA. As anticipated, this analysis indicated increased levels of Fibronectin (FN1), type I collagen (COL1A1), and Vimentin (VIM) in CCl_4_-treated mice in comparison to controls (**Figure 2G**). Notably, the CCl_4_ + cfAF mice displayed significantly lower levels of FN1 and COL1A1 when compared to those treated with CCl_4_ only (*P* < 0.01 by Student’s t-test), along with a modest decrease in VIM levels (**Figure 2G**). A comparison of these protein levels between CCl_4_ + cfAF mice and control mice revealed no significant difference (**Figure 2G**). Furthermore, higher magnification imaging illustrated reduced FN1 abundance and unremarkable liver architecture in CCl_4_ + cfAF mice relative to CCl_4_-treated mice (**Figure 2H)**. Therefore, cfAF treatment prevents CCl_4_-induced liver fibrosis by diminishing EMT and MFA.

### Treating human hepatic spheroids with cfAF prevents HSC activation

Hepatic spheroids composed of immortalized PH5CH2 human hepatocytes^30^ and LX2 human HSCs^29^ were utilized to assess whether cfAF treatment reduced human HSC activation *ex vivo*^51^. Spheroids were incubated with ethanol for 48 hours to model liver damage as previously described^52^, then either treated with 10 or 25% cfAF, or replaced with serum-free media (SFM; **Figure 3A**). Ethanol-free spheroids were used to assess baseline expression of EMT and MFA biomarkers in the absence of ethanol activation. We used immunostaining and quantitative analysis to measure COL1A1 protein levels in the hepatic spheroids and found that adding 25% cfAF to ethanol-treated spheroids resulted in a significant reduction of COL1A1 protein levels after 72h (*P* = 0.0261 by Student’s t-test; **Figure 3B, right and 3C**). The RNA levels of MFA biomarkers COL1A1, TGFΒR1, smooth muscle actin (ACTA2) were reduced after treatment with cfAF compared to SFM-only; the and the EMT biomarker VIM showed variable levels at different time points (**Figure 3D)**. Interestingly, ethanol-free (non-activated) spheroids exhibited elevated baseline expression of COL1A1 (**Figure 3B, left**). Treatment with 10% cfAF for 72 hours led to a reduction in COL1A1 expression, demonstrating a trend toward statistical significance (P = 0.06, Student’s t-test; **Figure 3C**). In conclusion, cfAF treatment inhibits HSC activation in a human hepatic spheroid model.

**Figure 3.**
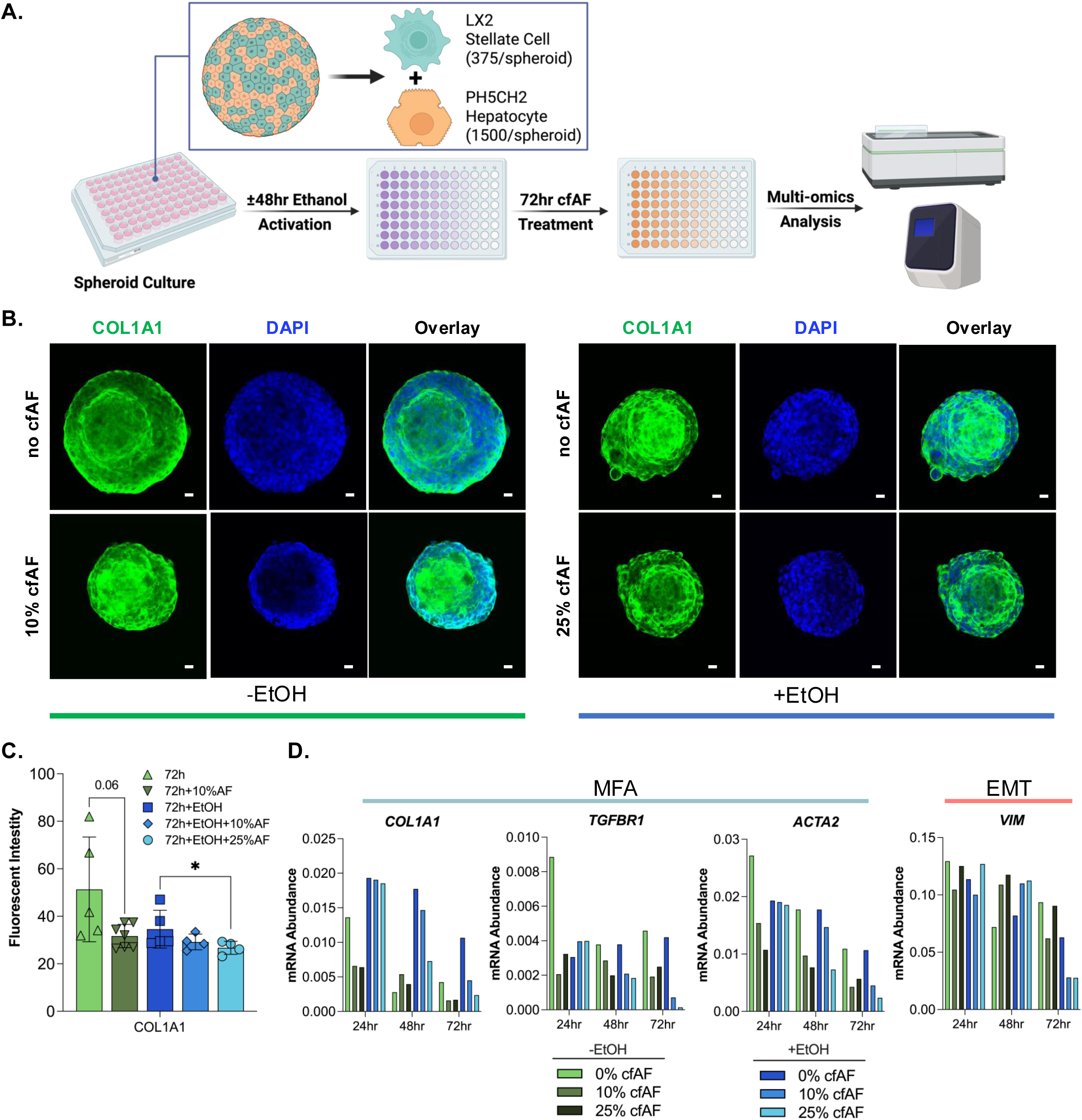
cfAF treatment reduces HSC activation in a human hepatic spheroid model. **A.** Schematic of spheroid culture and analysis. **B.** Representative images of ethanol-free hepatic spheroids (left) and ethanol-treated hepatic spheroids (right) cultured without (top) or with (bottom) cfAF and stained with anti-COL1A1 antibody and DAPI; scale bar = 20μm. **C.** Quantification of COL1A1 fluorescence intensity in non-activated (green bars) and activated (blue bars) spheroids relative to DAPI; N=5 for control or N=4 for cfAF-treated spheroids; \**P* < 0.05 by Student’s t-test. **D.** RNA was extracted from a minimum of eight individual hepatic spheroids—either activated (+ethanol, blue bars) or non-activated (−ethanol, green bars)—harvested at 24, 48, or 72 hours following treatment with 10% or 25% cfAF, or no treatment. RT-qPCR was performed to quantify mRNA abundance of MFA biomarkers and the EMT marker Vimentin (VIM), normalized to EEF1A1 expression.

### Cell-Free AF treatment prevents HSC migration and EMT-associated changes in RNA abundance

To determine whether cfAF can repress EMT *in vitro*, we first conducted the scratch test assay to measure the migration of LX2 or primary HSCs cultured with or without cfAF. Cell-free AF repressed cell migration in a dose-dependent manner (**Figure 4A and 4B).** We performed a growth curve analysis to evaluate if cell proliferation contributed to the reduction of the scratch area in any of the conditions, which can confound the migration assay. As expected, complete media supported robust cell proliferation, but LX2 HSCs cultured with cfAF proliferated more than those in SFM alone (**Figure 4C**). This indicates that cfAF promotes LX2 HSC proliferation compared to SFM, but this does not complicate the cell migration assay data interpretation where cfAF repressed migration. Transcript abundance measurements show that adding cfAF to SFM resulted in a dose-dependent decrease in EMT biomarkers (SNAIL, NCAD, FN1, VIM) compared to SFM alone (**Figure 4D**). Cell migration assays using primary human HSCs also indicate that cfAF represses their migration (**Figure 4E**) and reduces the levels of the EMT-related transcripts SLUG and SNAIL **(Figure 4F).** Therefore, cfAF represses EMT in human HSCs cultured *in vitro*.

**Figure 4.**
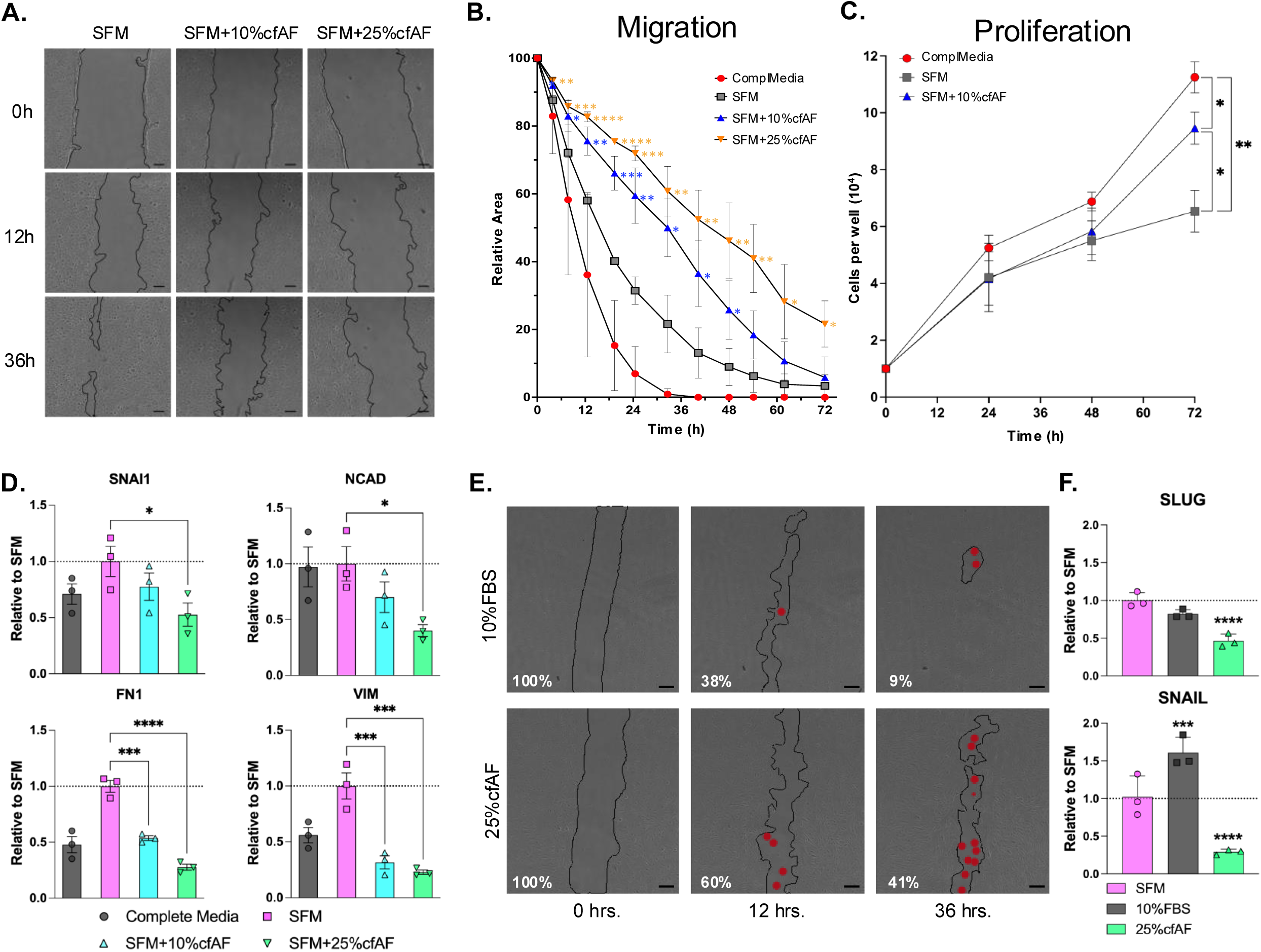
cfAF treatment represses EMT in HSCs. **A.** Representative phase contrast images of LX2 HSCs from the scratch test cell migration assay at 0-, 12-, and 36-hours post-scratch cultured in serum-free media (SFM), SFM + 10% cfAF, and SFM + 25% cfAF; scale bar = 200 μm. **B.** Quantification of the mean percentage of total starting scratch area (relative to time zero) over the 72-hour time course, calculated from three independent biological replicates; \**P* < 0.05, \*\**P* < 0.01, \*\*\**P* < 0.001, \*\*\*\**P* < 0.0001 by Two-way ANOVA with multiple comparisons. **C.** LX2 HSC growth curve over 72 hours showing mean cell number obtained each day from three independent biological replicates; “ComplMedia” refers to complete media (DMEM + 10% FBS); \**P* < 0.05, \*\**P* < 0.01 by Two-way ANOVA. **D.** RT-qPCR measurement of EMT biomarker transcript abundance from LX2 cell extracts following culture in complete media, SFM, or SFM with 10% or 25% cfAF; mean ddCt values were determined compared to GAPDH and data are shown as values relative to SFM from three independent biological replicates; \**P* < 0.05, \*\*\**P* < 0.001, \*\*\*\**P* < 0.0001 by One-way ANOVA. **E.** Phase contrast images of primary human HSCs from the scratch cell migration assay at 0-, 12-, and 36-hours post-scratch cultured in SFM + 10% FBS or SFM + 25% cfAF; scale bar = 200μm; red dots indicate regions within the scratched area that had one to several cells present. **F.** RT-qPCR measurement of SLUG or SNAIL RNA abundance from primary HSC extracts that had been cultured in SFM, SFM + 10% FBS, or SFM + 25% cfAF; mean ddCt values were calculated compared to GAPDH and data are shown as level relative to SFM from three independent biological replicates; \*\*\**P* < 0.001, \*\*\*\**P* < 0.0001 by One-way ANOVA with multi-comparisons.

### Cell-free AF treatment attenuates HSC myofibroblast activation in vitro

We next tested the hypothesis that cfAF treatment attenuates LX2 HSC myofibroblast activation *in vitro*. To limit experimental bias, we performed multi-omics analysis using RNA or protein extracted from LX2 HSCs cultured in serum-free media (SFM), SFM with 25% cfAF (+cfAF), SFM with TGFβ inhibitor A-83-01, SFM + TGFβ to stimulate HSC activation^53^, TGFβ-activated cells treated with A-83-01 as a positive control for attenuating activation^31^, or TGFβ-activated cells treated with cfAF (**Figure 5A**). We performed RNA sequencing and measured changes in RNA abundance by Deseq2 analysis^54^. This revealed thousands of changes in RNA abundance (**Figure 5B; Suppl Table S2**), including 1166 overlapping transcripts between TGFβ-activated LX2 HSCs and TGFβ-activated cells treated with A-83-01 or cfAF **(Figure 5C)**. This indicates that many transcripts altered upon LX2 HSCs activation also respond to adding A-83-01 or cfAF to TGFβ-activated cells **(Figure 5C).** To determine whether the 2345 TGFβ-responsive transcripts are altered oppositely by A-83-01 to attenuate activation or upon cfAF treatment, we compared their changes in TGFβ + A-83-01 or TGFβ + cfAF to TGFβ alone and performed linear regression analysis. As expected, we observe a strong negative correlation in the presence of A-83-01 (slope = -0.78, R^2^ = 0.72), consistent with it attenuating TGFβ-stimulated LX2 HSC activation **(Figure 5D).** Interestingly, we observe a similar trend when cfAF is added to TGFβ-activated LX2 HSCs (slope = -0.55, R^2^ = 0.53), indicating that cfAF can also dampen TGFβ-stimulated changes in global RNA levels **(Figure 5D).** We validated two key MFA-associated transcripts’ changes using RT-qPCR: COL1A1 and HGF are not only pro- or anti-inflammatory biomarkers in HSCs but are also drivers of either pro- or anti-inflammatory states in HSCs, respectively^55^. As expected, TGFβ activation increased COL1A1 abundance compared to SFM, but adding A-83-01 or cfAF to TGFβ-activated HSCs reduced COL1A1 RNA (**Figure 5E**, **top**). Consistent with a MFA of HSCs, hepatocyte growth factor (HGF)^55,56^ is lower in LX2 cells incubated with TGFβ, but is elevated upon adding either cfAF or A-83-01 to them (**Figure 5E, bottom**). Gene Set Enrichment Analysis (GSEA^57^) comparing the changes in RNA levels in SFM + TGFβ to SFM alone confirmed successful myofibroblast activation of LX2 HSCs: amongst the most highly enriched gene sets were TGFβ Signaling, Myc Targets, and EMT, while Interferon Alpha or Gamma Response, and IL6 JAK STAT3 Signaling were depleted **(Figure 5F).** We confirmed that A-83-01 indeed attenuates LX2 HSC activation, as GSEA revealed reduction in TGFβ Signaling, Myc Targets, and EMT and enrichment of Interferon Alpha Response, Interferon Gamma Response, and IL6 JAK STAT3 Signaling gene sets in TGFβ + A-83-01 vs TGFβ to TGFβ vs SFM **(Figure 5F).** Intriguingly, these were also observed among the most significantly altered gene sets when comparing transcript abundance changes between TGFβ + AF vs TGFβ wth TGFβ vs SFM **(Figure 5F).** Thus, treatment with either A-83-01 or cfAF potently represses transcript-level changes in gene sets associated with MFA in HSCs. Together, these data indicate that treating TGFβ-activated LX2 HSCs with cfAF attenuates myofibroblast activation-associated transcriptome changes.

**Figure 5.**
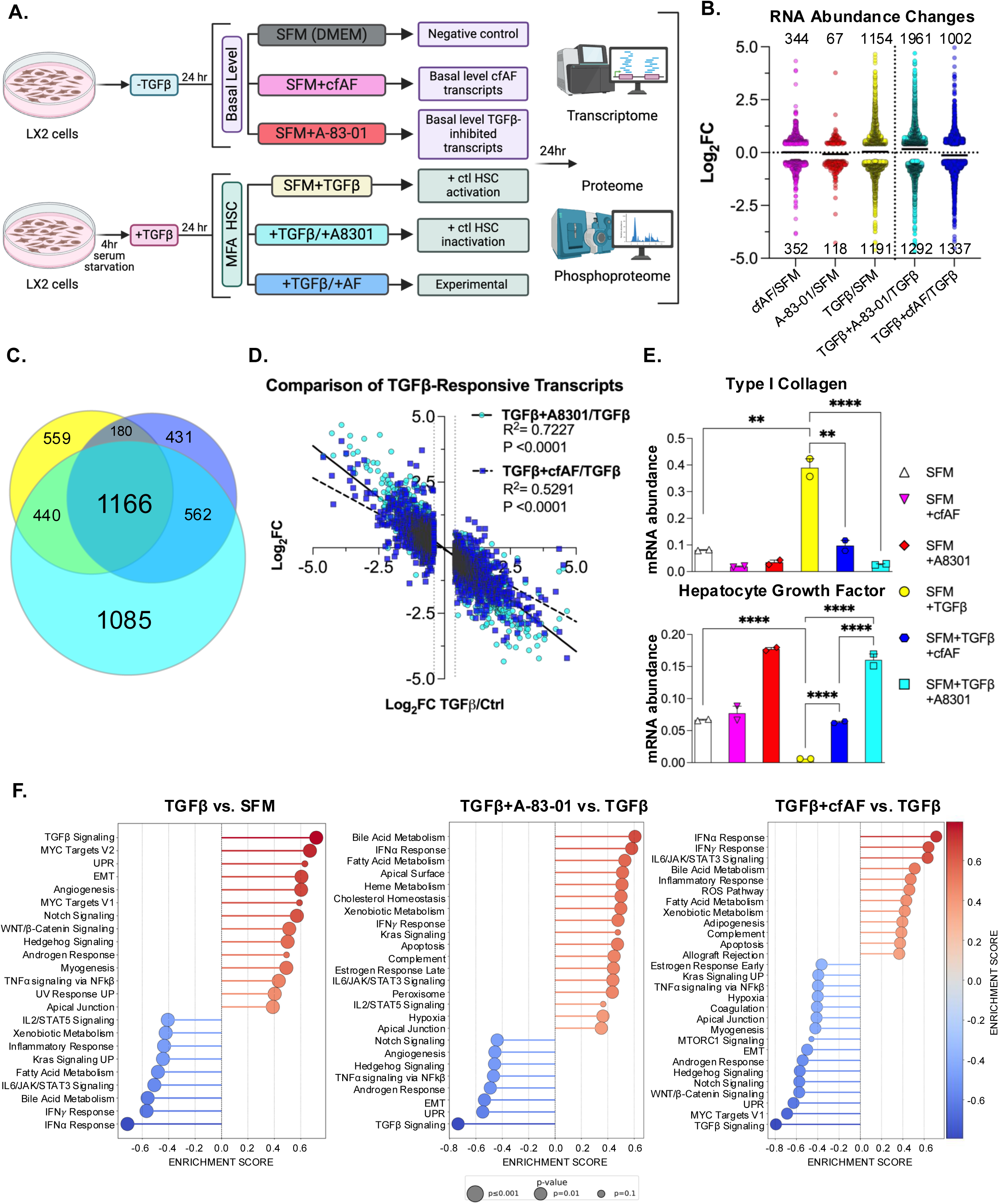
cfAF Treatment attenuates TGFβ-induced LX2 HSC activation-associated transcript level changes. A. Experimental approach schematic for LX2 HSC activation and analysis; each condition was performed using three independent biological replicate cell cultures. **B.** Log_2_ fold change values of changes in RNA abundance for extracts of LX2 HSCs that were cultured in 25% cfAF relative to SFM (cfAF/SFM), A-83-01/SFM, TGFβ/SFM, TGFβ + A-83-01/TGFβ, or TGFβ + 25% cfAF/TGFβ determined by Deseq2 analysis of RNA-seq data. **C.** Venn diagram illustrating the overlap of RNA abundance alterations as described in **B.** between TGFβ/SFM (yellow), TGFβ + A-83-01/TGFβ (teal), or TGFβ + cfAF/TGFβ (blue). **D.** Linear regression analysis of TGFβ-responsive transcripts (altered in TGFβ/SFM; log2 fold change on x-axis) compared to TGFβ + A-83-01/TGFβ (teal circles; log_2_ fold change on y-axis) or TGFβ + cfAF/TGFβ (blue squares; log_2_ fold change on y-axis). **E.** RT-qPCR analysis of RNA extracted from LX2 cells cultured as described in **B.** and measuring the mean mRNA abundance of COL1A1 or HGF compared to EEF1A1 from two independent biological replicates; \*\**P* < 0.01, \*\*\*\**P* < 0.0001 by One-way ANOVA with multi-comparisons**F.** Gene Set Enrichment Analysis (GSEA) from RNA-seq data analyzing TGFβ vs. SFM (left), TGFβ + A-83-01 vs TGFβ (middle), or TGFβ + cfAF vs TGFβ (right).

We measured global protein levels in the samples described above by LC-MS/MS to determine how protein abundance is altered **(Figure 6A** and **Suppl Table S3**) and how they overlap under these experimental conditions **(Figure 6B)**. Intriguingly, comparison of the proteome profiles of each culture condition shows two main clusters based on similarity: the first consists of TGFβ, cfAF + SFM, and SFM only, with the former being most dissimilar, and the second cluster consists of TGFβ + cfAF and TGFβ + A-83-01 **(Figure 6C)**. Therefore, TGFβ attenuation by A-83-01 or adding cfAF to TGFβ-activated LX2 HSCs show a similar proteomic profile. A notable protein (and RNA; see above) that increases during TGFβ-activation is COL1A1 **(Suppl Table S3)**, consistent with MFA and liver fibrosis. We validated this finding using antibody staining and immunofluorescence analysis in LX2 HSCs **(Figure 6D).** Type I Collagen levels were significantly reduced after adding A-83-01 to TGFβ-activated LX2 HSCs and to a lesser, but still significant, extent with cfAF treatment (**Figure 6D**; *P* < 0.0001 by One-way ANOVA). We also performed phosphoproteomics analysis on the same LX2 cell extracts. While transcriptome and proteome analysis revealed that the most changes observed were in A-83-01 + TGFβ compared to TGFβ alone, this approach found that the largest number of alterations were in cfAF + TGFβ compared to TGFβ (910 in the latter, 757 in the former, and fewer in the other comparisons; *P* < 0.05 and log2 fold-change ≥ |0.4**|; Figure 6E**, **Suppl Table S4**). Most of these were distinct to a certain treatment group comparison, but some overlap was observed, particularly when comparing the TGFβ-activated treatment with A-83-01 or cfAF (**Figure 6F**). The most significantly affected gene sets for alterations in phosphoprotein abundance between all comparisons were reduced in cfAF treatment of TGFβ-activated LX2 HSCs and included MAPK cascade and Wnt signaling pathway (**Figure 6G**). Interestingly, these were also reduced in CCl_4_-fibrotic mice that were treated with cfAF, compared to CCl_4_-treated alone at the RNA level (**Figure 2D**). Therefore, multiomics analysis reveals that cfAF can repress LX2 HSC myofibroblast activation via common and distinct gene expression and cell signaling alterations that are utilized by a direct TGFβ inhibitor.

**Figure 6.**
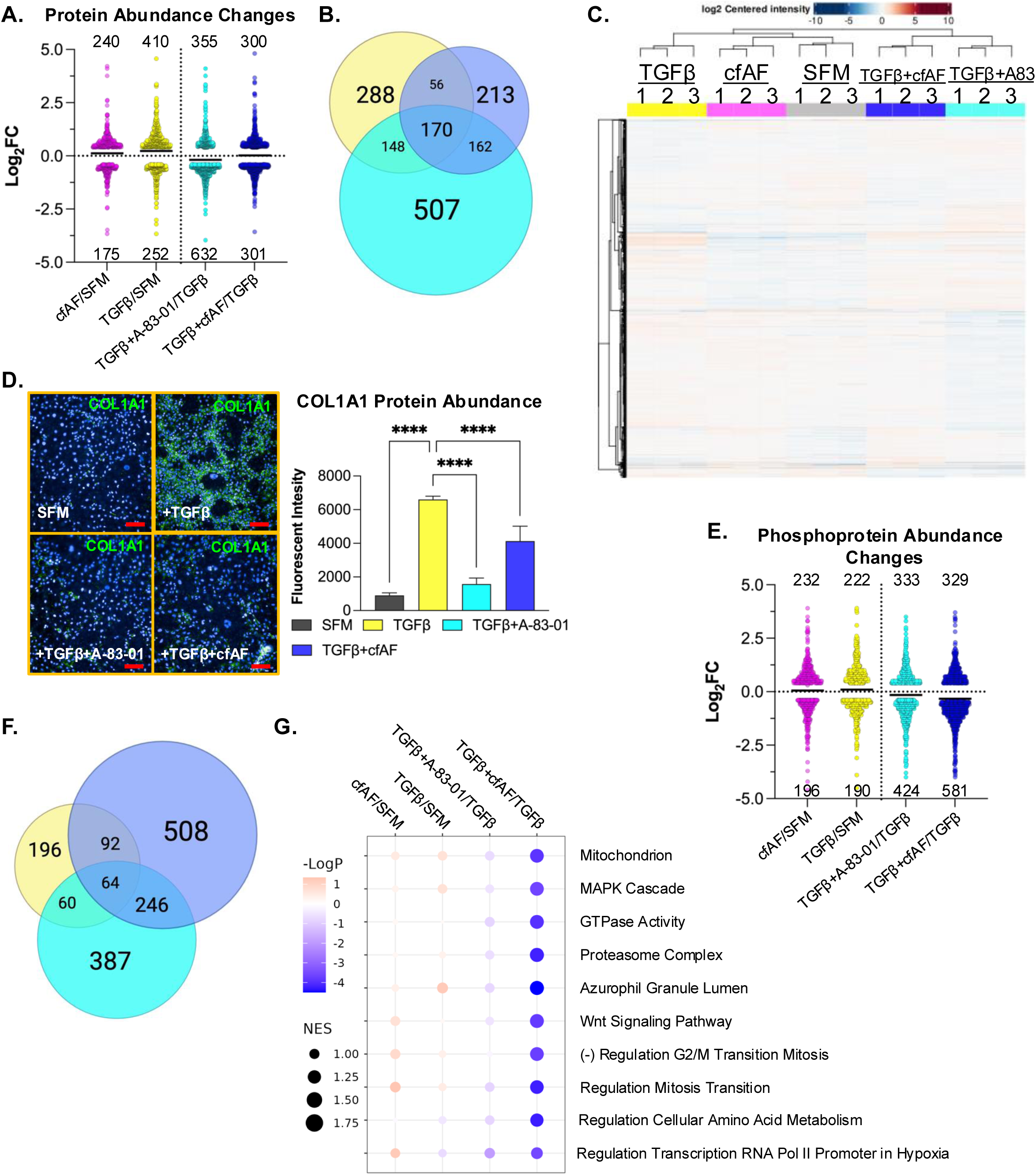
cfAF Treatment attenuates TGFβ-induced LX2 HSC activation-associated protein and phosphoprotein level changes. A. Log_2_ fold change values of changes in protein abundance measured by LC-MS/MS from extracts of LX2 HSCs that were cultured in 25% cfAF relative to SFM (cfAF/SFM), TGFβ/SFM, TGFβ + A-83-01/TGFβ, or TGFβ + 25% cfAF/TGFβ. **B.** Venn diagram illustrating the overlap of protein abundance alterations as described in **A.** between TGFβ/SFM (yellow), TGFβ + A-83-01/TGFβ (cyan), or TGFβ + cfAF/TGFβ (blue). **C.** Clustered heat map showing the top 500 most abundant proteins measured by LC-MS/MS for each replicate and each condition. **D.** Representative images of COL1A1 immunostaining in LX2 HSCs cultured in biological triplicate SFM, SFM + TGFβ, TGFβ + A-83-01, or TGFβ + 25% cfAF, collected and quantified on the Opera High Content Screening System and Harmony software with bar graph showing COL1A1 signal intensity relative to DAPI for each culture condition indicated; \*\*\*\**P* < 0.0001 by One-way ANOVA with multi-comparisons). Scale bars = 100 µm**. E.** Log_2_ fold change values of changes in phosphoprotein abundance measured by LC-MS/MS from extracts of LX2 HSCs as described in **A**. **F.** Venn diagram illustrating the overlap of phosphoprotein abundance alterations between TGFβ/SFM (yellow), TGFβ + A-83-01/TGFβ (cyan), and TGFβ + cfAF/TGFβ (blue). **G.** Gene Ontology Set Enrichment Analysis (GOSEA) of phosphoproteome data showing the top 10 categories with the lowest P-values and with the normalized enrichment score (NES) represented by dot size.

## Discussion

Here, we find that human cfAF treatment minimizes liver fibrosis and rescues failure-to-thrive in CCl_4_-treated mice without inducing a deleterious immune response (**Figure 1**). This protective effect is mediated by attenuating HSC myofibroblast activation and reducing the EMT (**Figures 2, 4, 5, 6**). We provide evidence strongly supporting this conclusion derived from the murine *in vivo* CCl_4_ model, a human hepatic spheroid *in vitro* model, and standard two-dimensional cell culture using both primary and immortalized human hepatic stellate cells. Our multi-omics approach limits experimental bias and facilitates discovery-based insight into the mechanisms through which cfAF elicits its therapeutic effects. To our knowledge, this is the first study of its type to administer human cfAF to mice via i.p. injection, analyze liver gene expression via transcriptomics and perform multi-omics analyses on human cells treated with cfAF. This provides unprecedented insight into cfAF’s therapeutic mechanisms and will serve as a resource for the scientific community. Hence, while our study suggests the exciting possibility that cfAF treatment could safely reduce or prevent liver fibrosis in humans, this also provides an accessible resource that can instruct interested parties on how cfAF influences gene expression and cell signaling.

### Fibrotic Liver Disease and cfAF Treatment

Liver fibrosis in humans is a highly variable clinical condition. It can be multi-etiological and a driver or symptom of different types of liver disease. In the case of environmentally induced liver fibrosis, such as excessive alcohol consumption, withdrawal of the insult does not cause immediate resolution of fibrosis, but rather, continued abstinence may allow for tissue remodeling and healing of the fibrotic liver scar tissue^58,59^. This gradual healing response is well-documented, but cases are rare. More often, behavior modification is unsuccessful, the liver is repeatedly injured, the fibrotic cascade is reinforced, and fibrosis worsens. At the cellular level, this manifests as: hepatocyte death and dysfunction, de-differentiation, and regenerative attempts; disruption of cholangiocyte function (e.g., altered bile acid metabolism); a shift in immune cell phenotypes with increased pro-inflammatory macrophages (e.g., activated Kupffer cells and monocyte-derived) changes in endothelial cells that reduce circulatory efficiency; and HSC myofibroblast activation and differentiation^4,58,60–62^. We show that cfAF protects CCl_4_-induced liver fibrosis and represses HSC myofibroblast activation. Several studies have shown that amniotic cell transplantation or treatment with conditioned media or extracellular vesicles (EVs) derived from amniotic cells can prevent liver fibrosis in rodent models^63,64^. Approximately 30-50 billion EVs per milliliter are observed in full-term cfAF^19,27,65,66^. These are derived from amnion epithelial cells (AECs), amniotic fluid cells (AFCs), and other fetal, maternal, and extra-embryonic sources and likely contribute to our observation that cfAF protects CCl_4_-induced liver fibrosis and represses HSC myofibroblast activation. Multi-omics analysis revealed remarkable overlap between *in vivo* **(Figure 2)** and *in vitro* (**Figures 4, 5, and 6**) models treated with cfAF, which indicates that it reduces the levels or activity of targets associated with EMT and MFA, and the signaling pathways Wnt, MAPK, mTOR, Notch/Hedgehog, and TNFa/NFkB. These are consistent with attenuated HSC activation, but the physiology of other liver cell types is likely influenced by cfAF treatment. Our fibrosis-focused transcriptome analysis of cfAF-treated mouse livers in the presence of CCl_4_ insult revealed increases in gene sets predominantly associated with immune function: M1 activation, type I and II interferon, Th1 and Th2 differentiation, and TLR signaling **(Figure 2).** This strongly suggests that cfAF alters the liver immune cell landscape. Amongst the downregulated sets, we observed HSC activation and liver fibrosis-associated signaling: myofibroblast regulation, EMT, Wnt, and TGFβ **(Figure 2).** Thus, cfAF treatment in this model activates immune surveillance while repressing HSC activation and signaling pathways deleterious to liver homeostasis. Although interpreting these results is complex, they may indicate the priming of a liver niche that is less permissive to hepatocellular carcinogenesis and that prevents extensive fibrosis. Further studies measuring cfAF’s effect on other liver cell types and in a hepatocellular carcinoma model would clarify these open questions.

### The Therapeutic Effects of cfAF

The regenerative biologics healthcare space has expanded in popularity recently, but the mechanisms through which many products’ therapeutic effects are mediated have not been explored. Until recently, cfAF was legally marketed and sold in the USA as a regenerative biologic, but in most cases, without data showing their efficacy or mechanism(s) of action. To confidently implement precision medicine and to explore the full potential of cfAF (and other regenerative biologics), efforts toward defining a comprehensive understanding of its components and how they elicit biological responses are critical. Full-term AF is a complex biofluid that contains various cell types and other insoluble components. Processing removes cells and debris larger than 0.2 µm but retains protein, nucleic acid, lipids/cholesterol, and other biomolecules. These can be either freely in solution or encapsulated in EVs. Interestingly, EVs from AECs or AFCs can also reduce liver fibrosis in the CCl_4_ model^63,64^. The diversity of biomolecules found in cfAF is likely much greater than in amniotic cell conditioned media or EVs, and the cost to produce cfAF compared to *in vitro* manufacturing of the latter is markedly lower. However, an open question regarding cfAF is whether a single fraction, component, or limited set of components is sufficient to convey a therapeutic effect in a certain disease/disorder-based context. Over millions of years, mammalian evolution has pressured developmental efficiency to yield the composition of AF that we currently observe. We speculate that various fractions of, and sets of biomolecules in, AF can elicit modulatory effects that could be desirable when repurposed as a regenerative biologic. Indeed, we discovered that AF EVs slightly activate EMT in fibroblasts and myoblasts, while total cfAF represses it, and EV-depleted cfAF has the strongest effect on reducing EMT^27^. Additional studies using more nuanced and sensitive fractionation techniques combined with comprehensive analyses of target cell/tissue types should begin to elucidate if there are specific “active fractions” of cfAF or if all parts of the whole are required to stimulate therapeutic or biological responses.

Here, we discover that cfAF treatment alters gene expression and signaling patterns in the mouse liver (**Figures 1 and 2**) or in human HSCs (**Figures 4, 5, and 6**). Consistent with our previous discoveries, cfAF potently represses cell migration and EMT-associated gene expression patterns (**Figure 4**). However, one of our major goals in this study was to limit our experimental bias and discover *de novo* mechanisms through which cfAF treatment alters cellular homeostasis. Interestingly, many changes in RNA abundance that occur upon TGFβ-mediated MFA of LX2 HSCs are reduced upon either A-83-01 or cfAF treatment (**Figure 5D**). Since A-83-01 blocks TGFβ signaling, it is unsurprising that its treatment of TGFβ-activated LX2 HSCs caused the largest number of RNA (**Figure 5B**) or protein **(Figure 6A)** abundance changes. In contrast, the largest number of changes in phosphoproteins was observed in the TGFβ + cfAF group (compared to TGFβ alone), where most of these were reduced (**Figure 6E**). Additionally, most of these were unique to this experimental group and did not overlap with the others (**Figure 6F**). This indicates that a large portion of the changes in RNA or protein abundance elicited by cfAF treatment of TGFβ-activated HSCs are like those stimulated by A-83-01 treatment but that cfAF treatment alters a unique and independent set of phosphoproteins. Thus, cfAF may have a generally repressive effect on kinase signaling that is manifest in reduced phosphoprotein abundance and prevents EMT and MFA in HSCs. In summary, our study reveals that cfAF prevents liver fibrosis by attenuating HSC activation and elucidates many of the therapeutic cellular-level effects that cfAF stimulates.

### Limitations

This study has several limitations that should be considered when interpreting the findings. First, the mechanism underlying the significant weight gain observed in mice treated with cfAF alone (**Figure 1B**) remains unclear. We hypothesize that the nutrient-rich composition of cfAF—particularly its abundance of proteins and lipids—may contribute to this effect; however, this remains speculative and requires further investigation.

Second, our hepatic spheroid culture system, designed to model the liver microenvironment in chronic liver disease, yielded variable and somewhat inconclusive results that did not fully align with the broader findings of this study. Although we followed previously published protocols (as described in the Materials and Methods), our data suggest that ethanol exposure did not reliably induce a fibrogenic or activated phenotype in the spheroids. Therefore, we cannot definitively conclude that ethanol served as an effective activating agent under our specific conditions. Notably, treatment with 10% cfAF led to reduced expression of activation-related biomarkers, such as COL1A1, even in the absence of ethanol, suggesting that cfAF may suppress basal levels of hepatic stellate cell activation. However, given the limitations of our spheroid culture system, including variability in response and technical constraints, further studies are warranted to clarify these effects and validate the observed trends.

## Conclusion

These data suggest that cfAF can reduce liver inflammation and fibrosis, primarily through the modulation of EMT, pro-fibrotic, and pro-inflammatory signaling pathways. These findings indicate that cfAF may serve as an effective treatment for liver inflammation and fibrosis induced by hepatotoxic damage. Moreover, cfAF appears to preserve liver function and protect against failure-to-thrive, suggesting its potential as a beneficial bridging agent for patients awaiting organ transplantation. However, further studies are needed to fully assess the potential of cfAF in human clinical trials for treating liver fibrosis and preventing cirrhosis or hepatocellular carcinoma (HCC).

## Acknowledgements

We thank the patients and their families for providing amniotic fluid to advance scientific discovery. We are grateful for their contribution to research. The authors want to acknowledge the UTMB ARC staff’s commitment and dedication to our murine colony husbandry. Additionally, the authors thank Meredith E. Weglarz from the UTMB Flow Cytometry Core for her unwavering support and contributions to this project. The authors also thank Omar Saldarriaga and Esteban Arroyave Sierra for their technical assistance using the Nanostring platform. We thank the UTMB Surgical Pathology Core for processing and preparing all tissue histology, the UTMB Obstetrics and Gynecology Department, and Merakris Therapeutics who graciously provided amniotic fluid. All cartoon images were created with licensed permission using BioRender.com.

## Availability of data and materials

The RNA-seq data has been deposited in GEO and can be accessed via GEO Series accession number GSE290015. The mass spectrometry proteomics data have been deposited to the ProteomeXchange Consortium via the PRIDE^67^ partner repository with the dataset identifier PXD060743 (and are accessible with token XXg5UVvaWtow until publicly available).

**Supplemental Table S1.** Nanostring Data

**Supplemental Table S2.** RNA sequencing data

**Supplemental Table S3.** Proteomics data

**Supplemental Table S4.** Phosphoproteomics data

**Supplemental Table S5.** Antibody information used for flow cytometry experiments.

**Supplemental Table S6.** Antibody information used for immunofluorescence experiments.

**Supplemental Table S7.** Primer sequences for PCR experiments.

## Graphical abstract

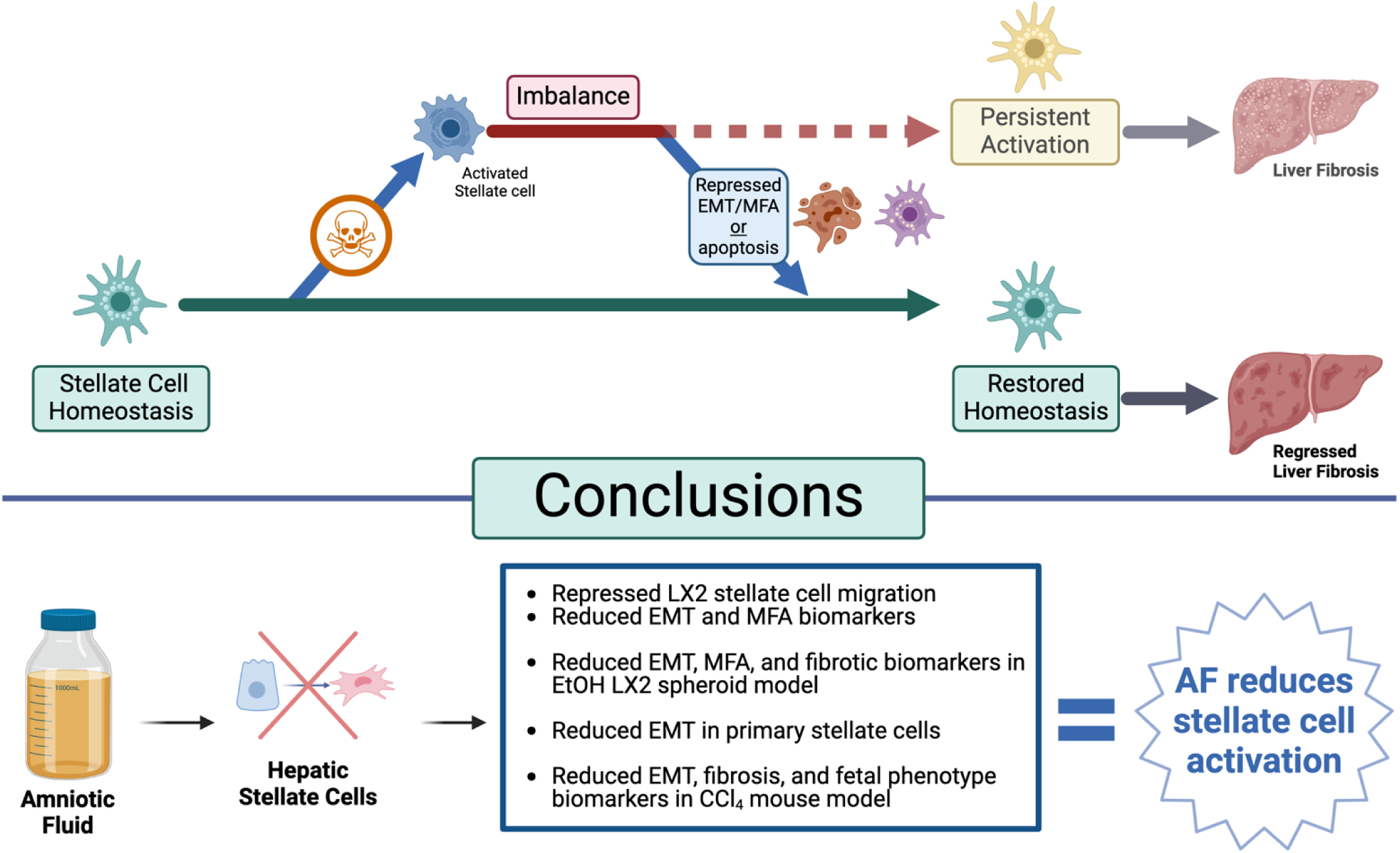

---

**Supplemental Figure 1.**
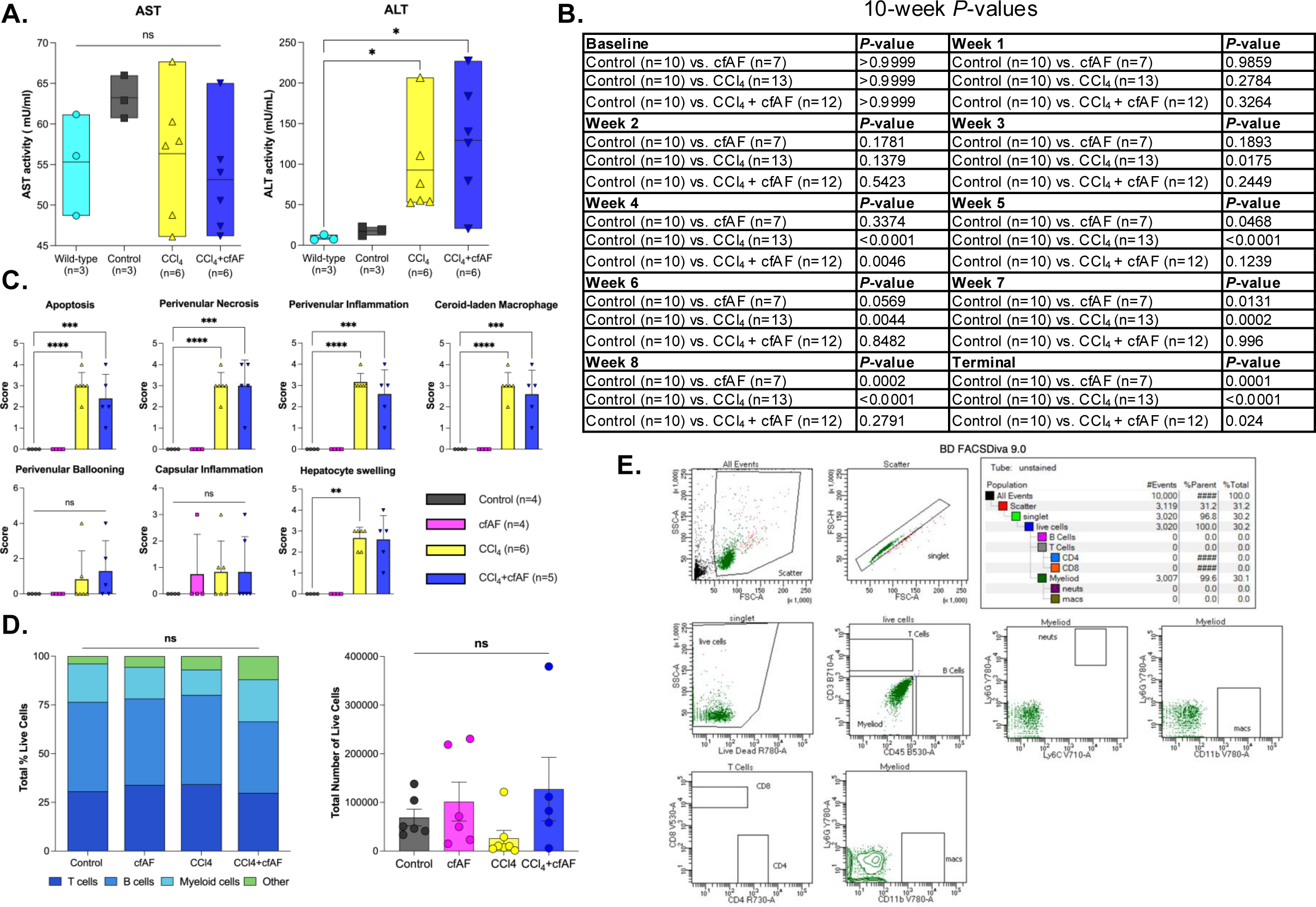
**A.** Liver function tests (AST and ALT) were performed on serum collected from mice at necropsy plotted as mean values (center line), with upper and lower bounds indicating the maximum and minimum values, respectively; N = 3 wild-type mice, N = 3 control mice, N = 6 CCl_4_ only mice, N = 6 CCl_4_+cfAF mice; \**P* < 0.05 by Student’s t-test. **B.** Results from Two-way ANOVA with multi-comparisons of mean mouse weight data for each group compared to control mice over 10-week CCl_4_ animal study (accompanies **Figure 1B**). **C**. Quantification of H&E-stained murine liver histology from the CCl_4_ animal study, scored blindly by an expert pathologist; N = 4 control mice, N = 4 cfAF only mice, N = 6 CCl_4_ only mice, N = 5 CCl_4_+cfAF mice; \*\**P* < 0.01, \*\*\**P* < 0.001, \*\*\*\**P* < 0.0001 by One-way ANOVA with multi-comparisons (accompanies **Figure 1C**). **D.** Flow cytometry analysis of PBMC T cells, B cells, myeloid cells, or other cells (**left**) or viable splenocyte abundance (**right**) across treatment groups; N = 6 control mice, N = 6 cfAF only mice, N = 7 CCl_4_ only mice, N = 5 CCl_4_+cfAF mice; statistical analysis by One-way ANOVA with multi-comparisons.

**Supplemental Figure 2.**
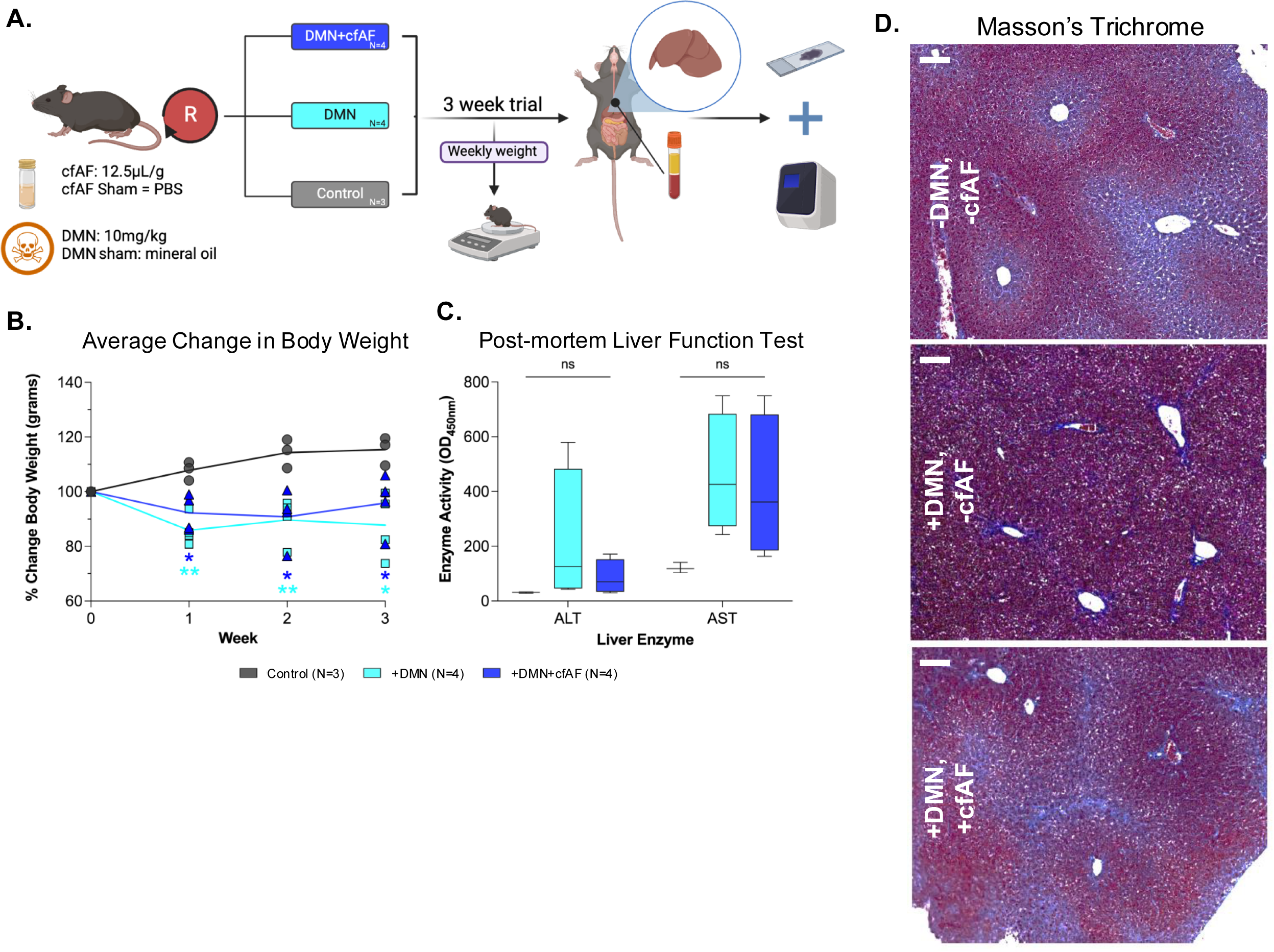
**A.** Overview of experimental design for modeling acute liver damage and cfAF administration in mice. **B.** Mean percent weight change by week, relative to starting body weight plotted by treatment group. N= 3 control mice, N = 4 DMN only mice, N = 4 DMN+cfAF mice; \**P* < 0.05, \*\**P* < 0.01 by Two-way ANOVA with multi-comparisons. **C.** Liver function tests (AST and ALT) performed on serum collected from mice at necropsy plotted as mean with upper and lower bounds corresponding to maximum and minimum values, respectively; N = 3 control mice, N = 4 DMN only mice, N = 4 DMN+cfAF mice; ns indicates not statistically significant by Student’s t-test. **D.** Liver sections from Control mice (top), DMN-treated mice (middle), and DMN-treated with cfAF mice (bottom) stained with Masson’s trichrome; shown at 20X magnification. Scale bars (upper left) = 100 µm.

**Supplemental Figure 3.**
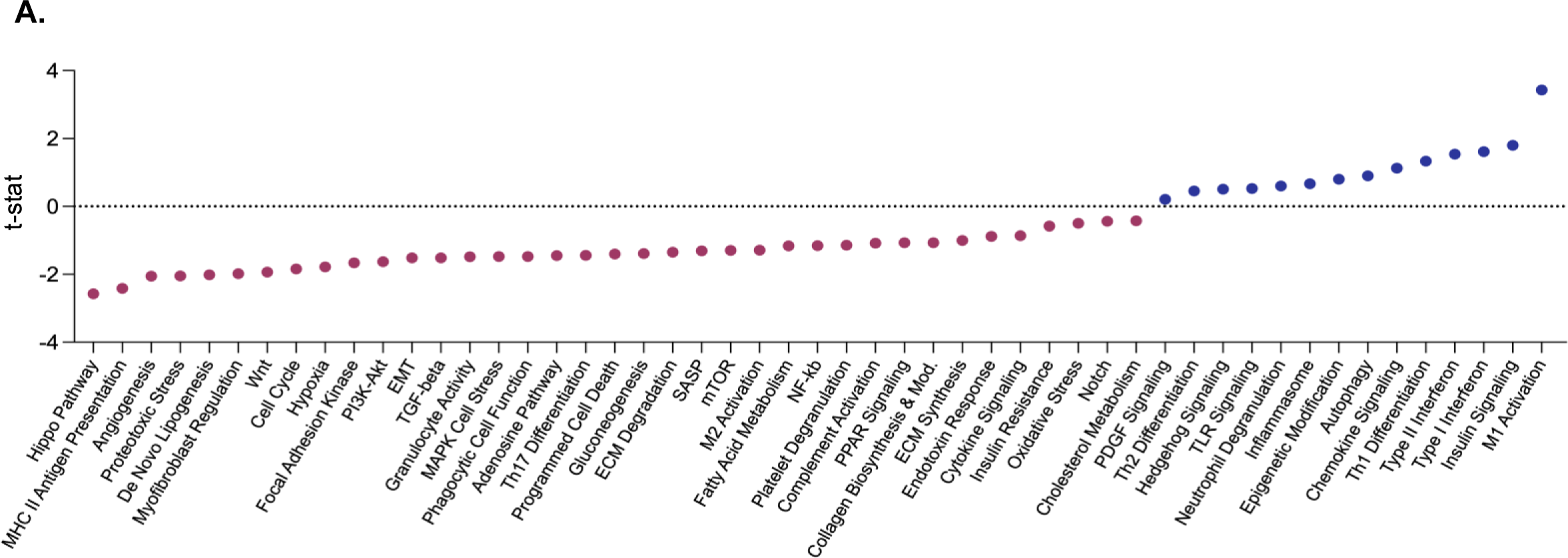
**A.** Complete pathway analysis of RNA level changes associated with enrichment (blue) or depletion (red) of pathways in CCl_4_ + cfAF mice versus CCl_4_ mice (accompanies **Figure 2E**).

## Graphical abstract

