## Supplemental Methods for "Amniotic Fluid Reduces Liver Fibrosis By Attenuating Hepatic Stellate Cell Activation"

**Supplemental (Full) Materials and Methods**

***Ethics statement.*** All patient samples (cfAF) and information were collected with written informed consent. Animal care and husbandry conformed to practices established by the Association for the Assessment and Accreditation of Laboratory Animal Care (AAALAC), The Guide for the Care and Use of Laboratory Animals, and the Animal Welfare Act. The Institutional Animal Care and Use Committee (IACUC) of The University of Texas Medical Branch approved all animal experiments in accordance with institutional guidelines (IACUC protocol #1706040B)*.*

***Animal Experiments.*** Wild-type female C57BL/6J mice between 6-8 weeks of age were purchased from Jackson Laboratory (Strain #:000664, Bar Harbor, ME). Mice were grouped based on test condition: sham control (n = 10), cfAF only (n = 7), CCl_4_ only (n = 13), or CCl_4_ plus cfAF (n = 12), for the CCl_4_ chronic liver fibrosis study; and sham control (n = 5), DMN only (n = 5), or DMN plus cfAF (n = 5) for the DMN acute liver damage study. Mice were treated twice weekly for nine weeks with CCl_4_ (2.25 µL/gram body weight or 10 mg/kg DMN) diluted 1:3 in food-grade olive oil or sham control (olive oil only). Either cfAF or saline control was injected i.p. twice weekly for nine weeks (on days when mice did not receive either DMN or CCl_4_) at a dose of 12.5 µL/gram body weight (stock cfAF [1 mg/mL]) or comparable volume of saline. Blood was collected using a capillary tube via the retroorbital approach under anesthesia at baseline, or at the terminal endpoint via direct cardiac puncture. Mice were weighed weekly to monitor health and fitness.

***Tissue collection.*** Mice were humanely euthanized by cervical dislocation, and a laparotomy incision was made from the sternal notch to the pubis. The liver was perfused *en bloc* by continuously injecting 5 mL of sterile Phosphate Buffered Saline (PBS) (Corning Life Sciences, Corning, NY) into the portal vein before tissue harvest. Samples for RNA and protein isolation were harvested from the left median lobe and then snap-frozen in liquid nitrogen (LN_2_). The left side of the median lobe was divided and half of it embedded in Tissue-Tek® O.C.T (Sakura, Tokyo, Japan) for immunohistology, and the other half in was placed in 10% formalin (Sigma-Aldrich, St. Louis, MO) for fixation prior to paraffin embedding. The spleen was also collected (see below) to analyze splenocytes.

***RNA and Protein Extraction from Liver Tissue.*** Ice cold RSB-100 lysis buffer (100 mM Tris-HCl pH 7.4, 0.5% NP-40, 0.5% Triton X-100, 0.1% SDS, 100 mM NaCl, and 2.5 mM MgCl_2_ + 0.1M PMSF and SIGMAFAST proteinase inhibitor cocktail [Cat#S8830]) was added to the snap-frozen tissue on ice, which was first homogenized manually using a syringe flange and subsequently by vortexing. The suspension was centrifuged at 13,000 rpm at 4°C, and the supernatant was collected and then equally divided for RNA isolation or storage of the protein extract. The RNA was purified using the ReliaPrep RNA Cell Miniprep System (Promega, Madison, WI) per the manufacturer’s protocol and quantified using a NanoDrop One™ spectrophotometer (Thermo Fisher Scientific, Waltham, MA). The protein did not require further purification and was quantified using a Bradford Assay (Bio-Rad Laboratories, Hercules, CA). Samples were stored at -80°C for downstream analyses.

***FFPE RNA Extraction.*** Formalin-fixed paraffin-embedded (FFPE) blocks and hematoxylin and eosin (H&E) slides were prepared by clinical pathology technicians in the histology core at UTMB. Tissue curls were cut from FFPE blocks at 50 micrometer (µm) using a microtome. RNA was extracted using the ReliaPrep FFPE Total RNA Miniprep System (Promega, Madison, WI) following the manufacturer’s protocol. RNA quantification was performed using a NanoDrop One™ spectrophotometer (Thermo Fisher Scientific, Waltham, MA) and Qubit™ Fluorometer 2.0 (Qubit, San Francisco, CA). RNA integrity was analyzed using the Tape Station RNA assay kit (Agilent Technologies, Santa Clara, CA).

***Liver Histology.*** 3 µm tissue sections were cut from FFPE blocks of mouse liver tissue and mounted onto glass microscopy slides. Hematoxylin and Eosin (H&E) and Masson’s trichrome stains were prepared by surgical pathology technicians at the UTMB Surgical Pathology Core and scored by an expert pathologist (H.S). The slides were scanned using an Aperio digital slide scanner (Leica Biosystems, Vista, CA) and quantified using ImageScope software (Leica Biosystems, Vista, CA).

***Splenocyte Harvest and Preparation.*** The spleen was aseptically harvested from each mouse and placed in a p60 dish containing 8-10mL of ice-cold RPMI/10% FBS. The spleen was placed between two glass slides and ruptured by rubbing the slides together until the organ was homogenized into small fragments. The homogenate was transferred back to the p60 dish. All contents were aspirated and centrifuged in a 15 mL conical tube at 453 x *g* for 5 min at 4°C. After removing the supernatant, the pellet was resuspended in 5 ml of RBC lysis solution (Qiagen, Germantown, MD) and incubated for 5 min at room temperature (RT). RBC lysis reactions were stopped by filling the 15 mL conical tube to the top with ice-cold PBS/10%FBS and centrifuged at 453 x *g* for 5 min at 4°C. The splenocytes were then resuspended in 1 mL of staining buffer (PBS + 1% FBS) in preparation for flow cytometry.

***Peripheral Blood Mononuclear Cell Preparation.*** Fresh whole blood from cardiac puncture was diluted 1:1 in PBS (Corning Life Sciences, Corning, NY) without calcium and magnesium and layered on top of 2 ml of Ficoll-Paque PLUS (Cytiva, Marlborough, MA) in a polystyrene flow cytometry tube. These were centrifuged at 1350 RPM for 30 minutes at 4°C using slow-brake deceleration on the centrifuge. The plasma layer of the gradient was collected and stored for future analyses. The buffy coat and histopaque layers were transferred to a 15 ml conical and diluted with 10 ml of PBS. The combined layers were then pelleted at 2000 RPM for 5 minutes at 4°C and transferred to a 1.5 ml snap cap tube. The cell suspension was washed twice with staining buffer (PBS + 1% FBS) and then further processed for flow cytometry analysis as noted below.

***Cell Preparation for Flow Cytometry and Flow Cytometry.*** Prior to staining and fixation, a small fraction (~100,000 cells) of splenocytes and PBMCs were reserved as unstained controls for proper gating and assessing autofluorescence/background signal levels. The remaining cells were prepared for staining. To evaluate cell viability, 1 µL of Zombie NIR™ amine-reactive dye (BioLegend, San Diego, CA) was added to the cells (100μL suspensions in staining buffer) and incubated at RT in the dark for 30 minutes, followed by a wash with 1 mL of cell staining buffer to remove excess dye. Cells were then resuspended in 100 µL staining buffer and stained with BV785 anti-mouse CD11b (BioLegend, 101243), PerCp/Cyanine5.5 anti-mouse CD3 (BioLegend, 100328), BV510 anti-mouse CD8a (BioLegend, 100752), Alexa700 anti-mouse CD4 (BioLegend, 100430), and FITC anti-mouse CD45R/B200 (BioLegend, 103206) antibodies and incubated for 20 min on ice in the dark. Stained splenocytes, PBMCs, and unstained controls were fixed with 4% paraformaldehyde (PFA) for 20 minutes on ice in the dark, washed, resuspended in staining buffer, and stored at 4°C in the dark. Antibody information is listed in **Suppl Table S5.** The splenocytes and PBMCs were analyzed using the BD LSR Fortessa Cell Analyzer under sterile conditions at the UTMB Flow Cytometry Core. Compensation beads (BioLegend, 424601) were prepared per the manufacturer’s guidelines prior to running the experimental samples. Experts in the core facility established all gating and compensation parameters. Results were analyzed by FlowJo Software version 10.8.1 (Tree Star, Inc., Ashland, OR, USA), and scatter plots showing gating parameters are provided in **Supplemental Figure 1E**

***Immunofluorescence.*** Antibody staining was performed on 15 or 20 µm liver sections (cut by cryo-microtome from O.C.T.-embedded liver tissue) that were mounted on frosted glass slides (Thermo Fisher Scientific, Waltham, MA). The samples were incubated in 100% ethanol for 20 minutes, followed by 1X PBS for 5 minutes. The slides were air dried, incubated in 0.1% Triton-X solution for 1 min, and washed three times with 1X PBS. Then, the slides were incubated with 0.3% Sudan Black in 70% ethanol in a beaker for 2 hours in the dark. The slides were then washed twice with 70% ethanol and then incubated in fresh sodium borohydride (1 mg/mL in PBS, pH 7.5) for 15 minutes. The slides were then washed twice with 1X PBS, and a DAKO pen was used to encircle the tissue on the slide. Upon drying, the slides were washed with ddH_2_O for 5 minutes. The slides were then treated with Proteinase K diluted in 1X TBS (1:1000 or 20 µg/mL) for 10 minutes at RT before washing in ddH_2_O for 3 minutes. The slides were then blocked overnight with 1X SBS (1% horse serum, 0.1g BSA, 0.1% fish gelatin, 50 mM EDTA) at 4°C in a humidity chamber. The following day, primary antibodies were added and incubated overnight at 4°C in a humidity chamber. Antibody information, including dilution and catalog number, are listed in **Supp Table S6.** The slides were then washed three times with 1X TBS, and the secondary antibodies were overlaid in 1X SBS and incubated for 1 hour at RT. After incubation, the slides were washed 5 times in TBS and then Prolong-Gold (Invitrogen) mounting media with DAPI was used to overlay the liver sections and glass coverslips were placed on top of them. Liver sections were imaged using a Nikon Eclipse Ti2 ZDrive A1 confocal microscope. Image analysis was performed using FIJI image analysis software version 1.0.

Cultured LX2 HSCs were fixed with 4% PFA in PBS for 15 minutes at RT, followed by three 5-minute washes with PBS. The cells were then blocked for 60 minutes using Blocking Buffer (CST #12411). The primary antibody was prepared in Antibody Dilution Buffer (CST #12378) at the manufacturer-recommended dilution. The primary antibody was added to the fixed HSCs and incubated overnight at 4°C, followed by three 5-minute washes with 1X PBS. The cells were then incubated with a fluorochrome-conjugated secondary antibody, diluted in Antibody Dilution Buffer (CST #12378), for 2 hours in the dark at RT. After three 5-minute PBS washes, the cells were counterstained with DAPI (0.5μg/mL). Imaging and analysis were performed using an Opera Phenix Plus High Content Screening System (PerkinElmer). Information for the antibodies used in immunofluorescence experiments is listed in **Supp Table S6**, which includes the manufacturer’s recommended dilution and clone information.

Spheroids were stained using previously established methods^28^ with the following changes. Briefly, spheroids were fixed using 4% PFA for 45 minutes. Following fixation, spheroids were resuspended in ethanol for storage prior to staining. When ready, spheroids were stained against COL1A1 as outlined above. After fixation and primary/secondary staining, spheroids were washed five times with cold PBS. Approximately 95% of the supernatant was removed during each of these washing steps. Spheroids were imaged at The University of Texas Medical Branch’s microscopy core using the Zeiss LSM 880 with Airyscan confocal microscope and ZEN Lite for image quantification and analysis.

***Nanostring Sprint RNA Measurement.*** The Nanostring v2 Murine Fibrosis panel was utilized (Nanostring Technologies, Seattle, WA) to measure the RNA abundance of genes associated with fibrosis. 150 ng of RNA isolated from FFPE liver tissue was prepared for the nCounter® SPRINT run following the manufacturer’s recommendations. The data analysis was performed using the Nanostring n-Solver Advanced Analysis 2.0 software utilizing R version 3.3.2. The cutoff used to determine statistically significant alterations in RNA abundance was *P* < 0.1.

***Cell Culture.*** LX2 HSCs were acquired from the Radhakrishnan Lab in the Department of Surgery at UTMB. PH5CH2 cells were acquired from Shelton Bradrick of the Garcia-Blanco Lab (formerly UTMB, now at UVA). Primary human HSCs were purchased from BioIVT (Westbury, NY). The cells were cultured in DMEM containing 10% FBS at 37°C in 5% CO_2_ and passaged upon reaching ~80-90% confluency as previously described^29,30^. LX2 HSCs were plated in a 12-well plate at 1.5 x 10^5^ cells/well and then cultured under the following conditions: complete media (DMEM + 10% FBS); Serum-free media, SFM (DMEM only); A-83-01 only (DMEM only + 1 μM A-83-01), cfAF only (DMEM only + 25% AF by volume). The selective TGFβ inhibitor, A-83-01 (Cat# 50-197-5873, STEMCELL Technologies, Vancouver, BC), was used as a positive control for TGFβ inhibition as previously described^31^. LX2 HSCs were serum starved for four hours and then treated with 10 ng/ml of TGFβ (Cat# ab50036, Abcam, Cambridge, UK) overnight to stimulate MFA. The following day the media were changed, and cells were then cultured with: TGFβ only (DMEM only + 10 ng/mL TGFβ), A-83-01 + TGFβ (DMEM only + 1 μM A-83-01+10 ng/mL TGFβ), or SFM + TGFβ + cfAF (DMEM only + 10 ng/mL TGFβ + 25% cfAF by volume). The cells were harvested and split equally by volume into: RSB-100 buffer (100 mM Tris-HCl pH 7.4, 0.5% NP-40, 0.5% Triton X-100, 0.1% SDS, 100 mM NaCl, and 2.5 mM MgCl2) to collect protein, or Trizol to collect RNA (see below), and stored at -20°C for RNA or protein abundance analyses. All cells utilized for experiments in this manuscript were authenticated using fingerprinting and tested for mycoplasma, annually.

**Primary Stellate Cell Culture and Maintenance**. Primary stellate cells were cultured in Stellate Cell Medium (SteCM; Cat#5301; Sciencell Research Laboratories, Carlsbad, CA) supplemented with Stellate Cell Growth Supplement (SteCGS; Cat #5352; Sciencell Research Laboratories, Carlsbad, CA) and 2% fetal bovine serum (FBS) to create a complete medium optimized for *in vitro* growth. Frozen stellate cells were thawed and plated following standard protocols previously mentioned, above. Briefly, 5 mL of cold SteCM was aliquoted into a sterile 15 mL conical tube. Cryovials were removed from liquid nitrogen storage and rapidly thawed in a 37°C water bath. Once thawed, vials were wiped dry, rinsed with 70% ethanol, and transferred to a sterile biosafety cabinet. The cell suspension was carefully transferred to the prepared conical tube using a pipette, and the cryovial was rinsed with 1 mL of cold SteCM, which was added to the tube. Cells were centrifuged at 250g for 5 minutes at 4°C, the supernatant was aspirated, and the pellet was resuspended in fresh medium. Cell viability and density were assessed using the Trypan Blue Exclusion Assay^32^. Cells were seeded at a density of 4,000 cells/cm² on collagen I-coated plates and allowed to adhere undisturbed for at least 12 hours. The medium was replaced the following day and subsequently changed every other day. Cells were sub-cultured upon reaching 90% confluency. The medium, 0.25% trypsin solution, and Dulbecco’s Phosphate Buffered Saline (DPBS, without calcium and magnesium) were pre-warmed to RT. The old medium was aspirated, cells were rinsed with DPBS, and trypsin was added to detach cells during a 2–5-minute incubation at 37°C. The detached cells were washed from the flask with an appropriate volume of medium, transferred to a centrifuge tube, and centrifuged at 250g for 5 minutes at 4°C. The supernatant was aspirated, and the pellet was resuspended in 1–2 mL of fresh medium. Cells were counted and re-seeded at 4,000 cells/cm² on collagen I-coated plates for continued culture.

***Hepatic Spheroid Culture.*** Immortalized hepatocytes (PH5CH2) and stellate cells (LX2) were cultured as described above. Once thawed or cultured respectively, cells were resuspended in spheroid media (William E Media w/ 1x pen/strep, 10ug/mL insulin, 5.5 μg/mL transferrin, 6.7 mg/mL sodium selenate, and 100nM dexamethasone) supplemented with 10% FBS. A small population of cells were then removed and counted using a hemocytometer with trypan blue staining, as previously mentioned. Corning round bottom ultralow attachment or Eliplasia 96 well plates were prefilled with 200 μl of spheroid media with 10% FBS. Then 1500 PH5CH2 hepatocyte cells and 375 LX2 HSCs were added to each well, and then carefully placed in an incubator for 5 days without disturbance to allow spheroid formation. Then spheroid coalescence was visually confirmed, then 50% of the volume of spheroid media was exchanged for serum-free spheroid media. After five days of 50% media replacement, spheroid culture conditions were considered essentially ‘Serum-Free’. Spheroid were cultured with 10% or 25% amniotic fluid in spheroid media or a control group that was given spheroid media alone. Spheroids were collected at 24, 48, or 72 hours after starting their ‘recovery’ and imaging or RNA extraction was performed (see below).

***Scratch Test Assay.*** LX2 cells were seeded as described above and allowed to grow to near confluence on a 6-well plate. The cells were then washed 3x with PBS and a small area was scratched off using a p200 pipet tip. Immediately prior to the ‘scratch,’ wells were subdivided into 4 conditions: Complete media (DMEM + 10% FBS), serum-free media (DMEM), and 10% or 25% cfAF (DMEM plus 10-25% cfAF respectively). Plates were monitored continuously using the CellAssist Automated Imaging system (Thrive Biotechnology, Boston MA).

***RNA extraction from cells/hepatic spheroids, reverse transcription, and qPCR***. RNA was extracted from cells, including spheroids, using Trizol-based methodology followed by the ReliaPrep RNA MiniPrep system (Promega), using the manufacturer's instructions and as previously described^33,34^. Two micrograms of purified RNA were reverse transcribed into cDNA using the iScript cDNA synthesis kit (Bio-Rad Laboratories, Hercules, CA). The resulting cDNA was diluted between 1:10 or 1:25 in nuclease-free water, and one microliter was used as the template in a 25-microliter qPCR in which PowerUP SYBR Green Master Mix (Thermo Fisher) master mix was used. Amplification and analysis were performed on a StepOne Applied Biosystems qPCR machine with StepOne software (Applied Biosystems, Waltham, MA) analyzed by the ΔΔCt method^35^. Product specificity was confirmed by melt curve analysis. All PCR experiments were performed in triplicate, and outliers (> 3 SDs) were excluded from the statistical analysis. Statistical calculations were performed using dCt (calculated relative to housekeeping gene as noted in figure legend) values while 2^-dCt^ values were plotted graphically to represent mRNA abundance. Primer sequences are provided in **Supp Table S7**.

***RNA sequencing.*** LX2 cell RNA was extracted as above, and the quality was measured by Agilent BioAnalyzer (all RIN values were > 8, indicating high quality). Libraries were prepared by polyA selection using the NEBNext Ultra II Directional kit, and RNA was sequenced on the Illumina NovaSeq X Plus. The data were analyzed as described by the ENCODE project^36^ and using the NFCore pipeline. Approximately 40 million paired-end reads were obtained for each sample replicate (n = 3 each) and mapped to the hg38 genome build with STAR^37^. Deseq2 (Bioconductor version 3.19) was used to measure changes in RNA abundance with the cutoff for significant changes set at: basemean > 100; p-adj < 0.01; log_2_ fold-change ≥ |0.4|. GSEA (v4.3.3) was performed using normalized read counts (TPM), and the significance cutoff for gene sets used was *P* < 0.1.

***Proteomics and Phosphoproteomics (LC-MS/MS)***. The samples were prepared following previously established methods^38^. Briefly, the agarose bead-bound proteins were washed several times with 50mM Triethylammonium bicarbonate (TEAB) pH 7.1, before being solubilized with 40μL of 5% SDS, 50 mM TEAB, pH 7.55 followed by a RT incubation for 30 minutes. The supernatant containing the proteins of interest was then transferred to a new tube, reduced by making the solution 10 mM Tris (2-carboxyethyl) phosphine (TCEP) (Thermo Fisher, #77720), and further incubated at 65°C for 10 minutes. The sample was then cooled to RT and 3.75 μL of 1 M iodoacetamide acid was added and allowed to react for 20 minutes in the dark after which 0.5 μL of 2 M DTT was added to quench the reaction. Then, 5 μL of 12% phosphoric acid was then added to the 50 μL protein solution followed by 350 μL of binding buffer (90% Methanol, 100 mM TEAB final; pH 7.1). The resulting solution was administered to an S-Trap spin column (Protifi, Farmingdale, NY) and passed through the column using a bench top centrifuge (30 second spin at 4,000 x *g*). The spin column was then washed three times with 400 μL of binding buffer and centrifuged (1200 RPM for 1 min). Trypsin (Promega, #V5280, Madison, WI) was then added to the protein mixture in a ratio of 1:25 in 50 mM TEAB, pH 8, and incubated at 37°C for 4 hours. Peptides were eluted with 80 μL of 50 mM TEAB, followed by 80 μL of 0.2% formic acid, and finally 80 μL of 50% acetonitrile, 0.2% formic acid. The resultant peptides were quantified by fluorometric peptide assay and peptide quantities were normalized. An aliquot was used for phoshopeptide enrichment using kit 88812 from Thermo. The other aliquot was dried in a speed vacuum (RT, 1.5 hours) and resuspended in 2% acetonitrile, 0.1% formic acid, 97.9% water and aliquoted into an autosampler vial.

***NanoLC MS/MS Analysis***. Peptide mixtures were analyzed by nanoflow liquid chromatography-tandem mass spectrometry (nanoLC-MS/MS) using a nano-LC chromatography system (UltiMate 3000 RSLCnano, Dionex), coupled on-line to a Thermo Orbitrap Eclipse mass spectrometer (Thermo Fisher Scientific, Waltham, MA) through a nanospray ion source.  A direct injection method is used onto an analytical column with a sonation column oven; Aurora (75 µm X 25 cm, 1.6 µm) from (ionopticks). After equilibrating the column in 98% solvent A (0.1% formic acid in water) and 2% solvent B (0.1% formic acid in acetonitrile (ACN)), the samples (2 µL in solvent A) were injected (300 nL/min) by gradient elution onto the C18 column as follows: isocratic at 2% B, 0-10 min; 2% to 27% 10-98 min, 27% to 45% B, 98-102 min; 45% to 90% B, 102-103 min; isocratic at 90% B, 103-104 min; 90% to 15%, 104-106 min; 15% to 90% 106-108 min; isocratic for two minutes; 90%-2%, 110-112 min; and isocratic at 2% B, till 120 min.

All LC-MS/MS data were acquired using an Orbitrap Eclipse in positive ion mode using a data-independent acquisition (DIA) method with a 16 Da windows from 400-1000 and a loop time of three seconds. The survey scans (m/z 350-1500) were acquired in the Orbitrap at 60,000 resolution (at m/z = 400) in centroid mode, with a maximum injection time of 118 msec and an AGC target of 100,000 ions. The S-lens RF level was set to 60. Isolation was performed in the quadrupole, and HCD MS/MS acquisition was performed in profile mode using the orbitrap at a resolution of 30000 using the following settings: collision energy = 33%, IT 54ms, AGC target = 50,000.

***Database Searching***

The raw data was demultiplexed to mzML using MSConvert. The resulting mzML files were searched in MSFragger using the recommended DIA-NN settings ([https://github.com/vdemichev/DiaNN](https://nam11.safelinks.protection.outlook.com/?url=https%3A%2F%2Fgithub.com%2Fvdemichev%2FDiaNN&data=05%7C02%7Cwirussel%40UTMB.EDU%7Ccb9f5de956544c3de93f08dc52782c16%7C7bef256d85db4526a72d31aea2546852%7C0%7C0%7C638475923290519684%7CUnknown%7CTWFpbGZsb3d8eyJWIjoiMC4wLjAwMDAiLCJQIjoiV2luMzIiLCJBTiI6Ik1haWwiLCJXVCI6Mn0%3D%7C0%7C%7C%7C&sdata=C7oze4tUuQUDT9kOJw7FsbOffcK%2Blf7qK6%2BljoPCW2w%3D&reserved=0)): Peptide length range 6-30, Precursor charge range 2-5, Precursor m/z range 300-1700, Fragment ion m/z range 200-2000, Protease set to Trypsin/P, 1 missed cleavage, 2 variable modifications, N-term excision on, C carbamidomethylation on, Ox(M) on, Precursor FDR (%) set to 1.0, Mass accuracy set to 20 ppm, MS1 accuracy set to 10 ppm, Use isotopologues and MBR turned on, Neural network classifier set to single-pass mode, Protein inference set to protein names from FASTA, Quantification strategy set to Robust LC (high accuracy), and Cross run-normalization set to RT-dependent. The files were searched against a database of human acquired from Uniprot (21^st^ January, 2022).

Statistical analysis was performed using Fragpipe-Analyst using an R script based on the ProteinGroup.txt file produced by DIA-NN^39,40^. First, contaminant proteins, reverse sequences and proteins identified “only by site” were filtered out. In addition, proteins that have been only identified by a single peptide and proteins not identified/quantified consistently in same condition are removed as well. The DIA data was converted to log_2_ scale, samples were grouped by conditions, and missing values were not imputed. Protein-wise linear models combined with empirical Bayes statistics were used for the differential expression analyses. The limma package from the R Bioconductor was used to generate a list of differentially expressed proteins for each pair-wise comparison. A cutoff of the adjusted p-value of 0.05 (Benjamini-Hochberg method) along with an absolute log_2_ fold change of |0.4| has been applied to determine significantly altered proteins in each pairwise comparison.

***Liver Function Test.*** Whole blood collected from terminal cardiac puncture was centrifuged in microtubes for 2-3 minutes at 2,000 RPM to collect serum. Serum samples were prepared for assessment of AST and ALT activity using assay kits ab105135 (AST) and ab105134 (ALT) per the manufacturer’s recommended protocol (Abcam, Cambridge, UK). Activity was analyzed using a plate reader at OD_450nm_ and converted to IU/L. All samples were run in technical triplicate, and statistical calculation was reported using a Student’s t-test.

***Amniotic Fluid Procurement and Processing*** As previously described^27^, non-smoker donors who tested negative for infectious disease and gestational diabetes who scheduled cesarean delivery at full term were considered eligible. The obstetrician collected the fluid using a red Robinson tube connected to a 60 mL syringe to carefully aspirate AF after hysterotomy was performed, and before delivery of the fetus.

The AF was stored on ice, transported to the research lab, and (using sterile technique in a class II biosafety cabinet for subsequent steps) was first passed through a 100 µm cell strainer (Falcon) to remove large debris and vernix. It was then centrifuged at 1200 RPM at 4°C to pellet cells. The supernatant was collected and centrifuged again at 4000 RPM for 15 min at 4°C and then several times at 8000 RPM for 30 min at 4°C to remove additional insoluble material. It was then further processed by passing through a resin-bonded glass fiber composite media 0.45 µm depth filter (Pall) for primary clarification, followed by a polyethersulfone 0.22 µm (Pall) sterilizing-grade filter. The Bradford assay was used to estimate each donor's protein concentration of prepared cfAF. At least three independent donors’ cfAF were then combined in equal concentration (working [1 mg/mL]) and stored in -20°C as 1mL aliquots.

***Statistical analysis*.** Comparisons between two experimental groups were performed with GraphPad Prism using Student’s unpaired t-test, and among more than two experimental groups were analyzed using either One-way ANOVA or Two-way ANOVA with Tukey’s post-hoc analysis for multiple comparisons. The data are expressed as mean values ± SD from three independent experiments or biological replicates unless otherwise indicated.
